# DiNovo: high-coverage, high-confidence *de novo* peptide sequencing using mirror proteases and deep learning

**DOI:** 10.1101/2025.03.20.643920

**Authors:** Zixuan Cao, Xueli Peng, Di Zhang, Piyu Zhou, Li Kang, Hao Chi, Ruitao Wu, Zhiyuan Cheng, Yao Zhang, Jiaxin Dai, Yanchang Li, Lijin Yao, Xinming Li, Jinghan Yang, Haipeng Wang, Ping Xu, Yan Fu

## Abstract

Despite the recent advancements driven by deep learning, *de novo* peptide sequencing is still constrained by incomplete peptide fragmentation and insufficient protein digestion in current single protease-based proteomic experiments. Here, we present a software system, named DiNovo, for high-coverage and confidence *de novo* peptide sequencing by leveraging the complementarity of mirror proteases. DiNovo is empowered by several innovative algorithms, including a mirror-spectra recognition algorithm independent of pre-sequencing, two sequencing algorithms based on deep learning and graph theory, respectively, and target-decoy mapping, a method for sequencing result evaluation free of prior peptide identification. Compared with the trypsin protease used alone, DiNovo using two pairs of mirror proteases led to two to three times high-confidence amino acids sequenced. Compared with previous single-protease *de novo* sequencing algorithms, DiNovo achieved much higher sequence coverages. DiNovo also showed great potential as a powerful complement or alternative to database search for peptide identification with quality control.

## Introduction

*De novo* peptide sequencing from tandem mass spectra imposes no prior restrictions on the space of possible amino acid sequences, thus allowing for the identification of peptides or proteins outside existing knowledge databases, e.g., neoantigens, noncanonical antigens, synthetic peptides, antibodies, venom proteins, metaproteomes, and proteins from unknown species or species without sequenced genomes [1–5]. Traditionally, *de novo* sequencing algorithms have predominantly used the graph representation of tandem mass spectra and dynamic programming to search for the optimal sequence, such as *PEAKS* [6], *PepNovo* [7], *pNovo* [8]. Later, the *Novor* [9] algorithm introduced a machine learning approach to predict amino acids. In recent years, with rapid advancements in deep learning, neural networks have been intensively applied to *de novo* sequencing, with *DeepNovo* [10,11] as the pioneer, and dozens of followers such as *pNovo3* [12], *PointNovo* [13], *Casanovo* [14,15], *PepNet* [16], *GraphNovo* [17], *Denovo-GCN* [18] and *π*-PrimeNovo [19]. These algorithms show great capabilities in learning the hidden features of spectra and improving the accuracy of *de novo* sequencing.

However, existing *de novo* sequencing methods are still facing many challenges. Firstly, incomplete peptide fragmentation results in poor coverage of fragment ions for many peptides, leading to inaccurate or incomplete sequencing results [20]. Secondly, the digestion efficiency of commonly used proteases, e.g., trypsin, is often insufficient, resulting in missed cleavages of proteins, so that undigestible peptides cannot be sequenced. Thirdly, although recent deep learning-based algorithms, like GraphNovo [17], can fill in missing ions to some extent through trained network, these predicted ions still lack experimental evidence, making it difficult to confirm the correctness of the predicted sequences. Furthermore, the evaluation of *de novo* sequencing algorithms heavily depends on constructing benchmark datasets from peptide-spectrum matches (PSMs) obtained by database search, which impedes the direct performance comparison between the two approaches to peptide identification.

Mirror protease technology is a powerful way to improve fragment ion coverage of tandem mass spectra [21]. It typically uses two proteases to cleave proteins at the C- and N-termini of the same specific amino acid(s), such as K/R for trypsin and LysargiNase (Ac-LysargiNase), producing mirror peptides that share identical intermediate sequences but have the specific amino acid at the C- and N-termini, respectively. Such mirror peptides exhibit complementarity in their mass spectra, known as mirror spectra. Fragment ions absent in one spectrum can be present in the other spectrum [22]. For example, in higher-energy collisional dissociation (HCD) mass spectra, trypsin-digested peptides usually have rich *y*-type fragment ions, while LysargiNase-digested peptides have rich *b*-type fragment ions. Although *de novo* sequencing methods based on mirror proteases, such as *pNovoM* [23] and *Lys-Sequencer* [24], have been developed, they still suffer from several serious flaws. Firstly, there is still a lack of available software to support the entire workflow of mirror-spectra recognition and peptide sequencing. For instance, Lys-Sequencer has not been made public at all. Meanwhile, pNovoM lacks the program of mirror-spectra recognition, a crucial prerequisite for mirror peptide sequencing, in its publicly available package. Secondly, existing methods use only one pair of mirror proteases, which leaves the incomplete-digestion problem unsettled, leading to many sequences undetectable. Thirdly, some key algorithmic problems have not been adequately addressed. For example, recognizing which spectral pairs are mirror spectra is the first step in using mirror spectra for *de novo* sequencing. In pNovoM, the recognition of mirror spectra relies on pre-sequencing of peptides from separate spectra obtained by single-protease digestion, which is both time-consuming and sensitive to spectrum quality, and the results of recognition lack an effective quality control standard. Moreover, deep learning techniques have not yet been explored for *de novo* sequencing of mirror peptides. All these issues hinder the practical application of the mirror-protease strategy in proteomics.

In this paper, we present ***DiNovo***, a comprehensive software system that supports the full workflow of *de novo* peptide sequencing from tandem mass spectra generated by mirror proteases. The features of DiNovo include support for multiple pairs of mirror proteases, a novel mirror-spectra recognition algorithm independent of pre-sequencing and with quality control, two optional *de novo* sequencing algorithms based on deep learning and graph theory, respectively, and a target-decoy approach for confidence evaluation of *de novo* sequencing results in the presence of sequence database but free of prior peptide identification. We evaluated the performance of DiNovo using two pairs of mirror proteases including trypsin/LysargiNase, and Lys-C/Lys-N, which were used to digest proteins from *E. coli* and yeast proteomes. By taking full advantage of the complementarity of mirror spectra, DiNovo achieved much higher sequence coverage and confidence than state-of-the-art single-protease *de novo* sequencing algorithms. Compared with the trypsin protease used alone, DiNovo using two pairs of mirror proteases led to two to three times high-confidence amino acids sequenced. Furthermore, DiNovo identified a comparable number of proteins to database search at the same FDR level, showing great potential as a powerful complement or even an alternative to database search for protein identification. DiNovo is designed to dramatically increase the coverage and confidence of *de novo* peptide sequencing and to facilitate mirror-protease proteomics.

## Results

### DiNovo workflow

Figure 1 outlines the workflow and modules of DiNovo. In the first step, protein samples are digested by one or more pairs of mirror proteases, and the resulting peptides are analyzed using liquid chromatography coupled with tandem mass spectrometry (LC-MS/MS). In this work, two pairs of mirror proteases are used, including trypsin/LysargiNase, and Lys-C/Lys-N. To be specific, trypsin/LysargiNase cleaves C-/N-terminal to lysine and arginine, while Lys-C/Lys-N cleaves C-/N-terminal to lysine alone, producing longer peptides.

**Fig. 1.**
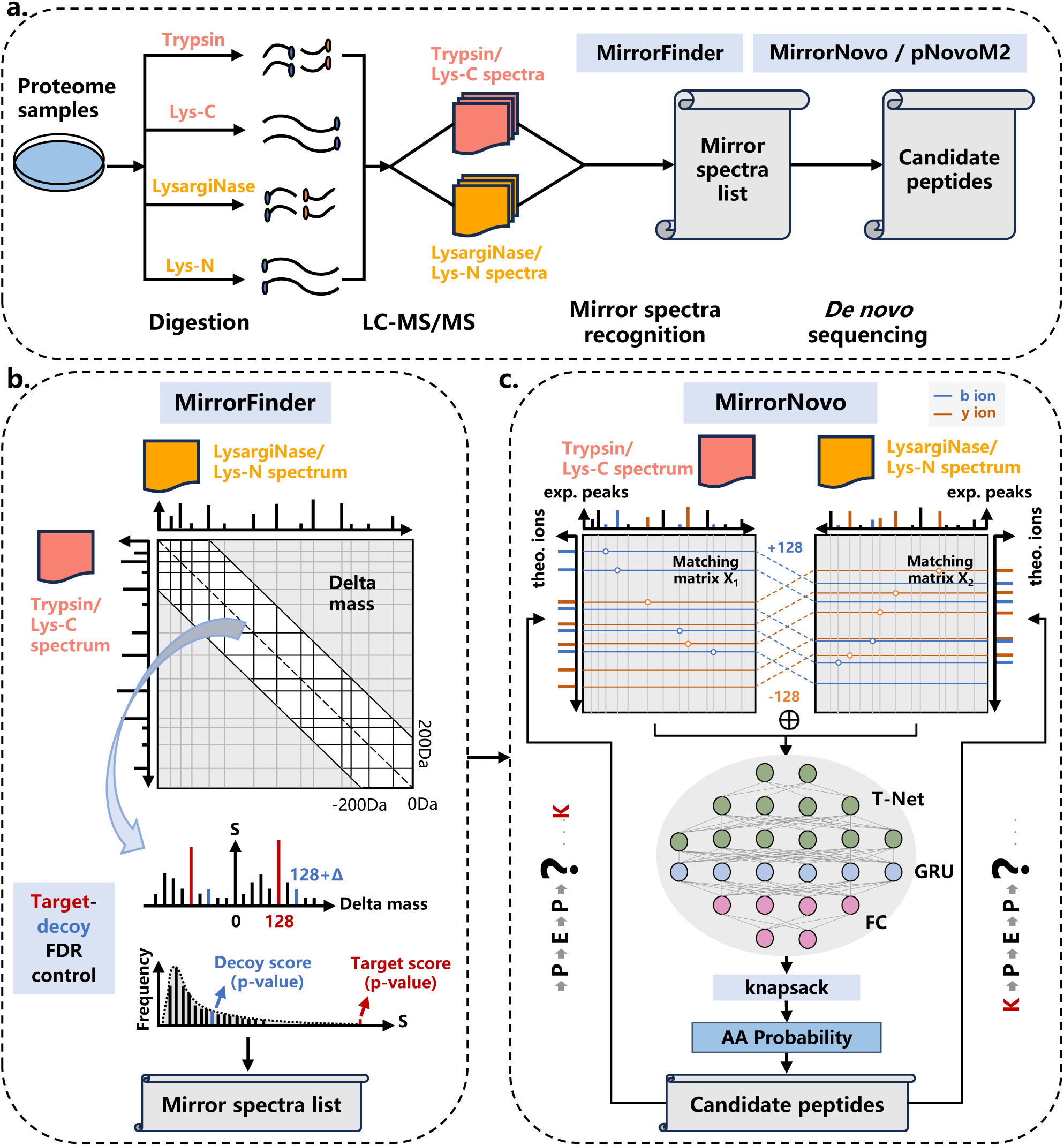
Algorithmic workflow and modules of DiNovo. **(a)** The workflow of DiNovo software. Proteome samples are first digested using multiple mirror proteases, followed by LC-MS/MS analysis to obtain tandem mass spectra. Then, mirror spectral pairs are recognized by MirrorFinder algorithm and peptides are sequenced by MirrorNovo or pNovoM2 algorithm. **(b)** MirrorFinder, the mirror-spectra recognition module of DiNovo, which leverages the distribution of fragment ion mass differences (delta masses) between two spectra to recognize mirror spectra and employs a target-decoy strategy for recognition error control. **(c)** MirrorNovo, the prime *de novo* sequencing module of DiNovo, which makes use of a designed neural network architecture to capture the peak features of mirror spectra and predict the probabilities of amino acids.

In the second step, mirror spectral pairs are recognized using a novel algorithm, called *MirrorFinder*. This algorithm does not rely on pre-sequencing of peptides from separate spectra, making it less sensitive to spectrum quality and more efficient. It directly utilizes the internal information of spectra to match spectral pairs, and employs a matching score to measure the possibility of a pair of spectra being mirror spectra (Figure 1b, Methods). The internal information includes precursor mass, fragment ion mass, and fragment ion intensity, which possess specific patterns for mirror spectral pairs. Furthermore, a target-decoy strategy is used to estimate and control the false discovery rate (FDR) of mirror spectral pairs. More details about the MirrorFinder algorithm are given in the Methods section.

In the third step, peptides are sequenced from the recognized mirror spectra using two developed algorithms. The first algorithm, called *MirrorNovo*, is based on a deep neural network (DNN), as shown in Figure 1c. Specifically, a T-Net is used to extract peak features from the feature matrix for matching peaks and theoretical fragment ions. Then, a Gated Recurrent Unit (GRU) layer is employed to capture the relationships between adjacent peak features. Finally, a fully connected (FC) layer outputs the probabilities of amino acids. In addition, a knapsack algorithm is applied at each step to constrain the search space. The second sequencing algorithm is *pNovoM2* (Figure S10), an improved version of pNovoM [23], which is based on graph representation of spectra and dynamic programming. While MirrorNovo is more accurate than pNovoM2, it requires GPU(s) for acceptable running speed. On the other hand, pNovoM2 runs on a CPU and is much faster than MirrorNovo when GPU(s) are unavailable. We recommend using MirrorNovo if GPU(s) are available, and pNovoM2 otherwise. In addition to mirror spectra, all separate spectra are also subjected to *de novo* sequencing for as high sequence coverage as possible. Further details on the sequencing algorithms can be found in the Methods section.

### Performance evaluation

To evaluate the performance of DiNovo, we constructed eight datasets using *E. coli* and yeast proteome samples, which were digested by two pairs of mirror proteases, i.e., trypsin/LysargiNase and Lys-C/Lys-N (see Methods for details on sample preparation and mass spectrometry experiments). We compared our mirror-protease strategy with the conventional single-protease strategy for *de novo* sequencing. We also compared DiNovo with previous single-protease *de novo* sequencing algorithms as well as the commonly used database search approach for peptide identification. Additionally, we evaluated the accuracy and speed of the two built-in sequencing algorithms in DiNovo, i.e., MirrorNovo and pNovoM2.

The traditional approach to evaluating *de novo* sequencing results primarily relies on constructing a benchmark dataset from peptide identifications obtained by protein sequence database search. However, this method cannot guarantee data quality due to the inherent error rate in database search results. Moreover, it directly prevents the horizontal performance comparison between *de novo* sequencing and database search. Another approach is to map *de novo* peptides to protein sequences in the database, and evaluate the number of peptide sequences mapped [16]. However, this approach lacks error control, as incorrect sequences can randomly match database proteins.

In this paper, we propose *target-decoy (TD) mapping* to evaluate *de novo* sequencing results, which is free of database search and allows for error control (Figure 2a, Methods). Specifically, we map all *de novo* peptides to a target-decoy protein database and count the number of target or decoy matches for FDR estimation, just as in the TD database search approach [25,26].

**Fig. 2.**
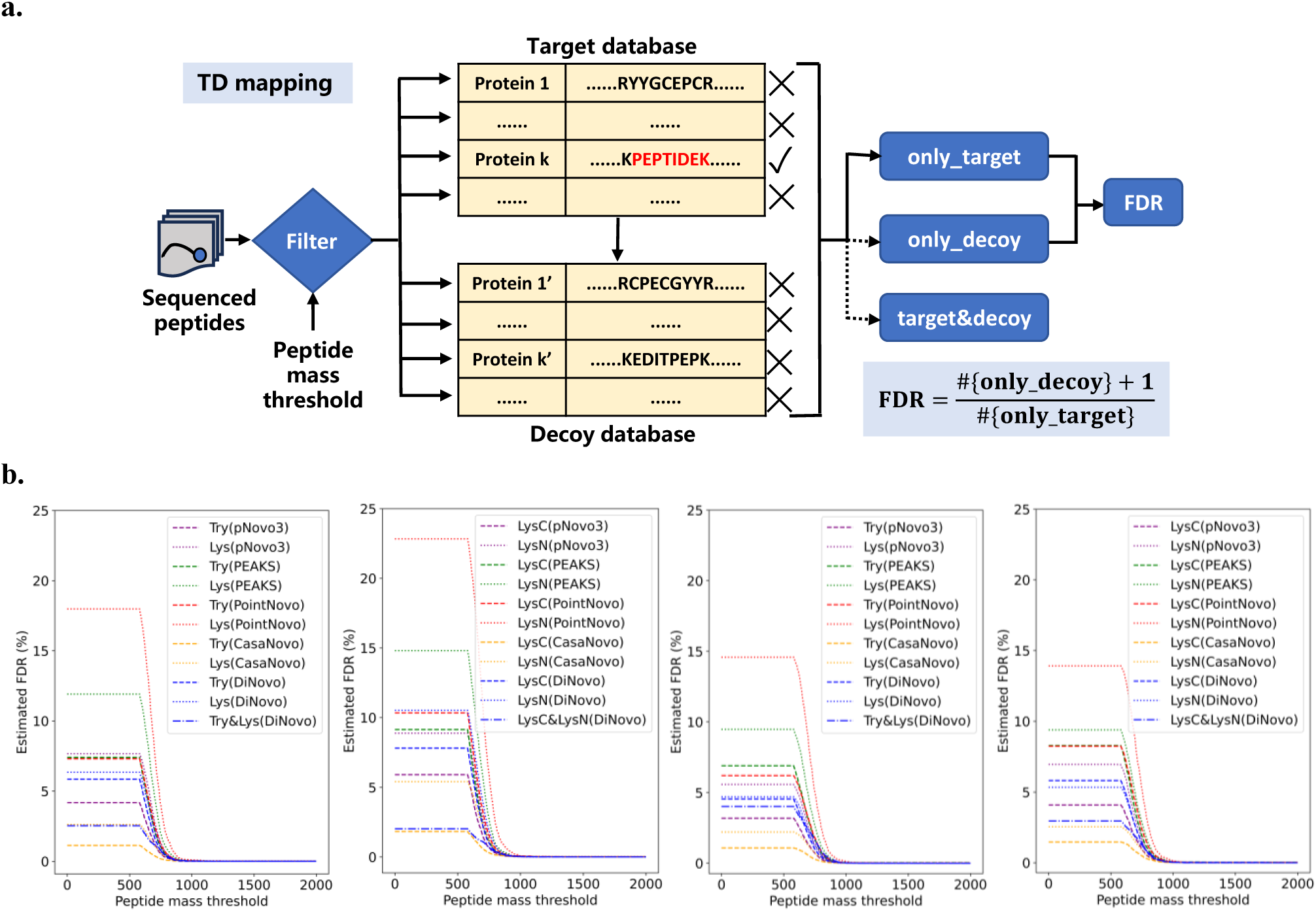
Quality control of *de novo* sequencing results. **(a)** The schematic diagram of TD mapping method. The decoy database is generated by reversing or shuffling the sequences in the original (target) database. Sequenced peptides whose masses above a certain threshold are mapped to the protein sequences in both the target and decoy databases, and the FDR is subsequently estimated based on the mapping results. **(b)** FDR estimated by TD mapping as a function of peptide mass threshold on the datasets of *E. coli* (first two) and yeast (last two) for various sequencing algorithms. Try and Lys are abbreviations of trypsin and LysargiNase, respectively.

Considering that short peptides are more likely to produce random matches, we filter the *de novo* peptides according to their precursor masses, which provide a more continuous measure than peptide lengths. Figure 2b shows how the estimated FDR changes with the precursor mass threshold. It is evident that the FDR curve declines sharply as the mass threshold increases, and stabilizes at close to zero after reaching 700-800 Da. As the mass threshold increases, mismatched peptides are gradually excluded. We ultimately chose a mass threshold corresponding to 1% FDR for each dataset. The threshold slightly differs between datasets (Table S1).

Furthermore, we focus on high-confidence sequencing results. A *de novo* peptide is considered high-confidence if it achieves 100% ion coverage, meaning that every amino acid residue of the peptide is supported by at least one spectral peak in the mass spectrum (considering only the top 200 spectral peaks). These high-confidence peptides are backed by more complete experimental evidence and are more convincing in practical applications.

### Near-complete fragment ion coverage in mirror spectra

The absence of some fragment ions presents a major challenge for *de novo* peptide sequencing. Our strategy in this paper is to improve the ion coverage using mirror proteases. To demonstrate the effectiveness of this strategy, we calculated the ion coverage of separate spectra and mirror spectra recognized by MirrorFinder. Figure 3 and S1 show that mirror spectra significantly complemented the missing ions in separate spectra on both *E. coli* and yeast datasets, which is highly beneficial for subsequent *de novo* peptide sequencing. Notably, a large portion of mirror spectra achieved near-complete fragment ion coverage, with average coverages of 98.4% for the *E. coli* datasets and 98% for the yeast datasets. In contrast, the average ion coverages for the separate spectra were only 90.2% and 89.7% for the *E. coli* and yeast datasets, respectively.

**Fig. 3.**
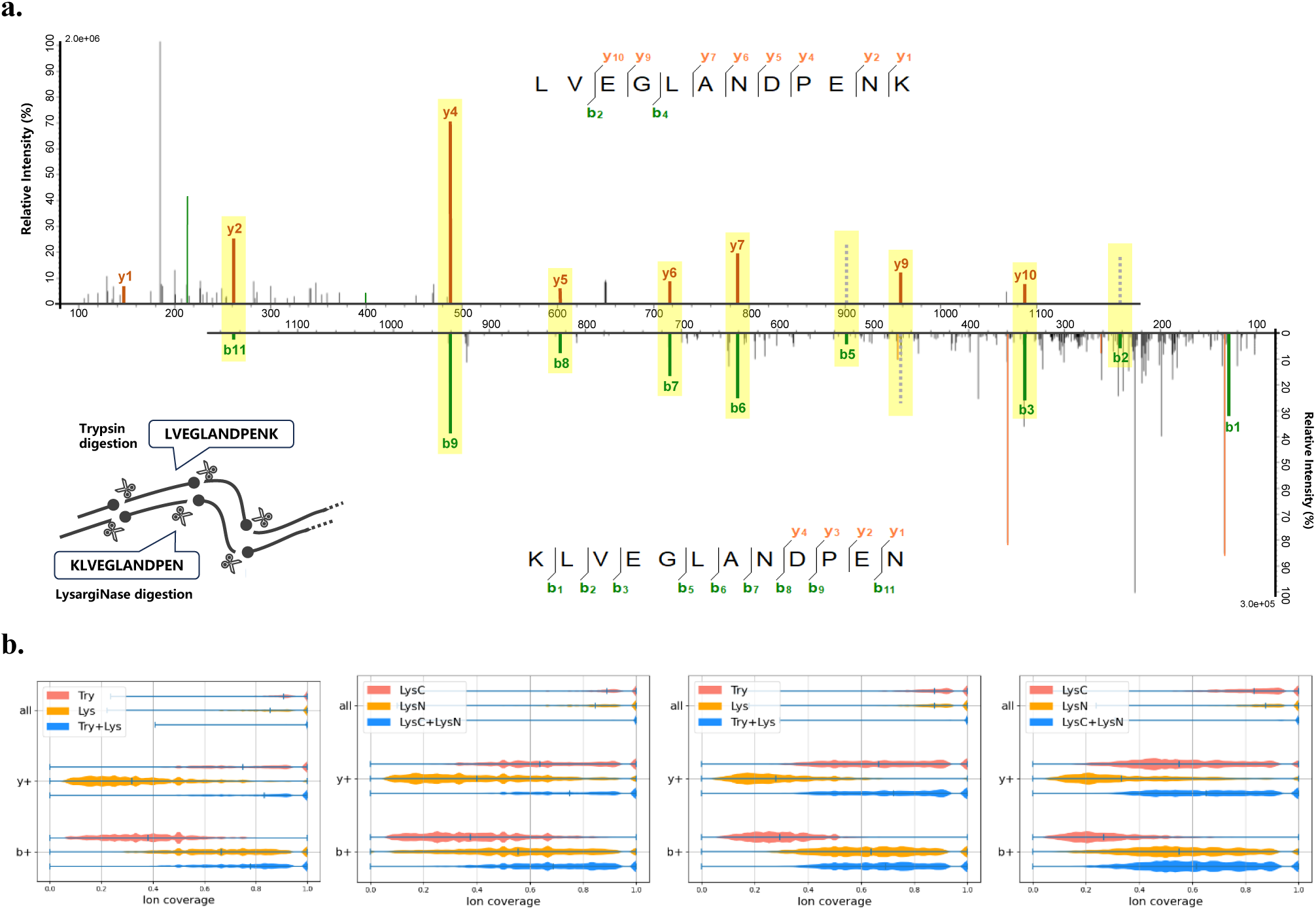
Near-complete fragment ion coverage due to mirror-protease strategy. **(a)** Illustration of the mirror-protease strategy and the complementarity of mirror spectra. The mirror proteases trypsin and LysargiNase cleave proteins at the C- and N-termini of lysines and arginines, respectively. The complementary fragment ions are highlighted by the yellow bar. Fragment ion absent in one spectrum are mostly present in the other, which improves the coverage of fragment ions. **(b)** Fragment ion coverages on the datasets of *E. coli* (first two) and yeast (last two). The *y*+ (*b*+) ion coverage is calculated as the proportion of theoretical *y*+ (*b*+) ions with matching spectral peaks. In mirror spectra, the spectral peak is matched once one of the mirror spectra is matched. “All” ion coverage is the proportion of fragmentation sites identified. A fragmentation site is considered identified when any of the *b*+, *b*++ (if any), *y*+, *y*++ (if any), or *a*+ ions corresponding to that site is matched.

### Comparison with single-protease strategy

To compare the mirror-protease strategy with the commonly used single-protease strategy for *de novo* peptide sequencing, we used DiNovo and PEAKS, with the latter being the industrial standard software for *de novo* sequencing, to sequence the *E. coli* and yeast proteomes, which were digested by trypsin/LysargiNase and Lys-C/Lys-N. We evaluated the number of peptides sequenced at an estimated 1% FDR, as well as amino acid coverage (the proportion of sequenced amino acids in the database), and protein coverage (the proportion of identified proteins). Throughout this paper, MirrorNovo was used within DiNovo unless otherwise specified.

As shown by Figure 4a and 4b, the mirror-protease strategy (DiNovo) significantly outperformed the single-protease strategy (PEAKS) in terms of all three metrics. Using either pair of mirror proteases, Try+Lys or LysC+LysN, was superior to using any of the four proteases alone. Combined use of the two pairs of mirror proteases within DiNovo resulted in the highest sequencing coverage and confidence. On the combined results of four proteases (All) for the *E. coli* (yeast) datasets, DiNovo sequenced 150.4% (154.8%) more high-confidence peptides than PEAKS, with 53.3% (57.6%) higher amino acid coverage and 16.4% (14.3%) higher protein coverage. Compared to the traditional single-protease strategy, such as Try(PEAKS), the advantages of DiNovo using two pairs of mirror proteases were more pronounced, as evidenced by sequencing 447% (573.6%) more high-confidence peptides for the *E. coli* (yeast) datasets, corresponding to 154.8% (226%) higher amino acid coverage and 279.5% (394.8%) higher protein coverage. Notably, DiNovo with one pair of mirror proteases (Try+Lys) surpassed the combined results of PEAKS with four proteases (All), revealing the immense contribution of mirror spectra to peptide sequencing. Furthermore, DiNovo identified 68.2% (68.1%) of *E. coli* (yeast) high-confidence proteins in the database, demonstrating the capability of multiple mirror-protease strategy in protein identification. The advantage of DiNovo over PEAKS was greater in terms of the number of peptides than amino acid and protein coverages. This was because DiNovo sequenced many mirror peptides with large sequence redundancy.

**Fig. 4.**
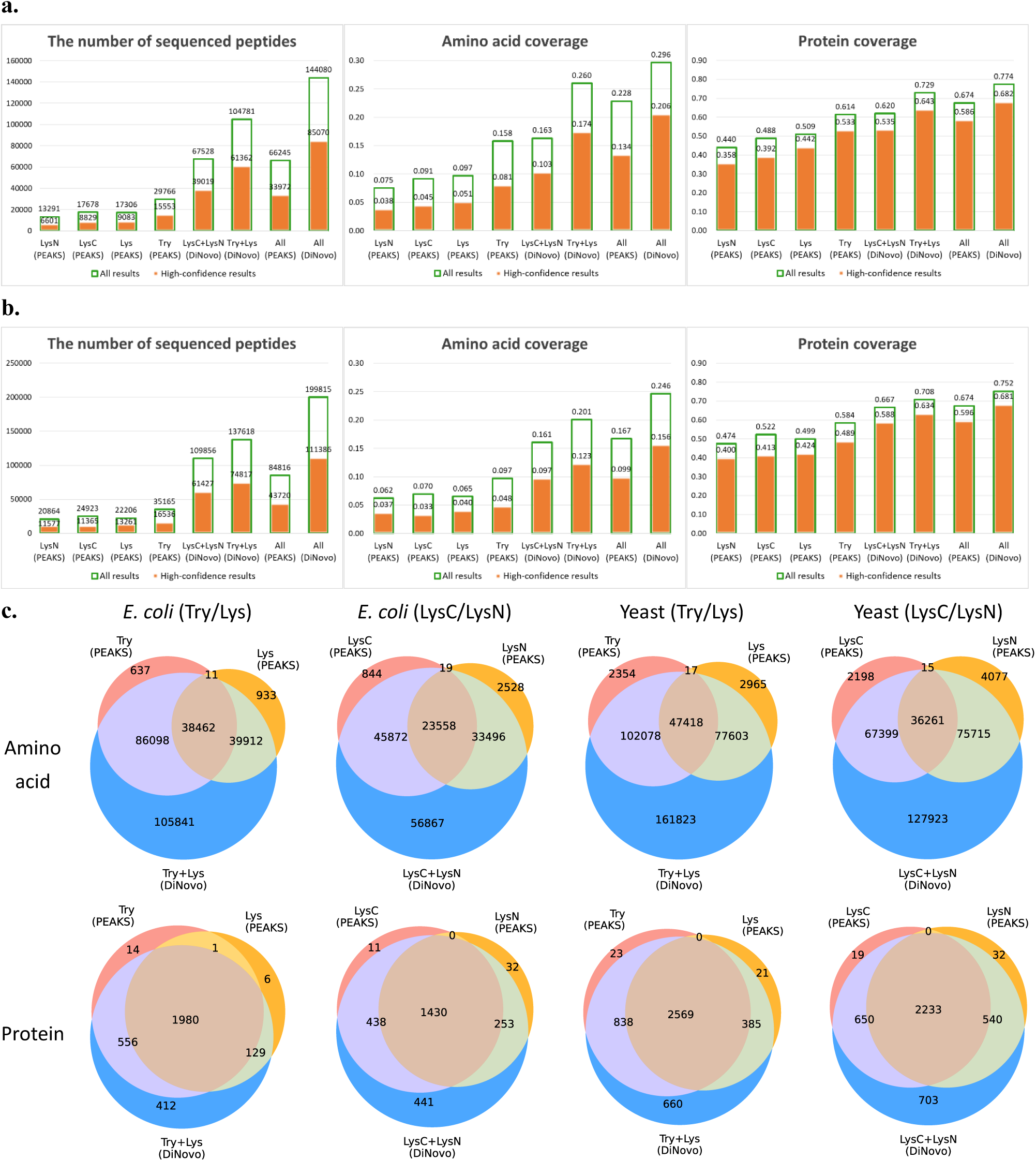
Comparison of *de novo* sequencing results obtained by single-protease and mirror-protease strategies. **(a)** *De novo* sequencing results on the *E. coli* datasets. **(b)** *De novo* sequencing results on the yeast datasets. Single-protease spectra were analyzed by PEAKS, and mirror spectra were analyzed by DiNovo. All(PEAKS) represents the union of LysN(PEAKS), LysC(PEAKS), Lys(PEAKS) and Try(PEAKS), and All(DiNovo) for the union of LysC+LysN(DiNovo) and Try+Lys(DiNovo). Green hollow bars indicate all results, while orange solid bars indicate high-confidence (full fragment ion coverage) results. **(c)** Venn diagrams of high-confidence amino acids and proteins sequenced by DiNovo and PEAKS.

Figure 4c further shows the intersection of high-confidence amino acids and proteins sequenced by DiNovo and PEAKS. Across all datasets with four proteases, DiNovo covered 96.6% to 99% of the high-confidence amino acids and 98% to 99.2% of the high-confidence proteins sequenced by PEAKS, while sequencing 56,867 to 161,823 additional high-confidence amino acids and 412 to 703 additional high-confidence proteins. More results are given in Supplementary Information (Figure S2, S3).

### Comparison with different *de novo* sequencing algorithms

To further demonstrate the advantages of mirror-protease strategy in *de novo* peptide sequencing, we compared the performance of DiNovo with more single-protease *de novo* sequencing algorithms. The description and configuration of these various software tools are detailed in Methods 4.3. The comparison was conducted on the combined results of four proteases for the *E. coli* and yeast datasets. As in the previous section, sequenced peptides were mapped to the target-decoy database, and the estimated FDR for each algorithm was controlled at 1% using different mass thresholds (Table S1). We report high-confidence results below.

The results in Figure 5a and 5b show that DiNovo outperformed all competitors at the peptide, amino acid, and protein levels for both *E. coli* and yeast datasets. Specifically, DiNovo sequenced 129.2% to 188% more *E. coli* peptides and 110.2% to 194.2% more yeast peptides than other algorithms, including Casanovo**, which was trained on an extremely large dataset. In terms of amino acid coverage, DiNovo exceeded Casanovo** by 42.8% (36%) for the *E. coli* (yeast) datasets and surpassed other algorithms by 51.7% to 73.6%. At the protein level, DiNovo possessed 11.6% to 19.5% higher protein coverage than other algorithms. Figure 5c shows the significant overlap of high-confidence amino acids sequenced by DiNovo and other algorithms. DiNovo covered most (96.7%-99.2%) amino acids of other algorithms.

**Fig. 5.**
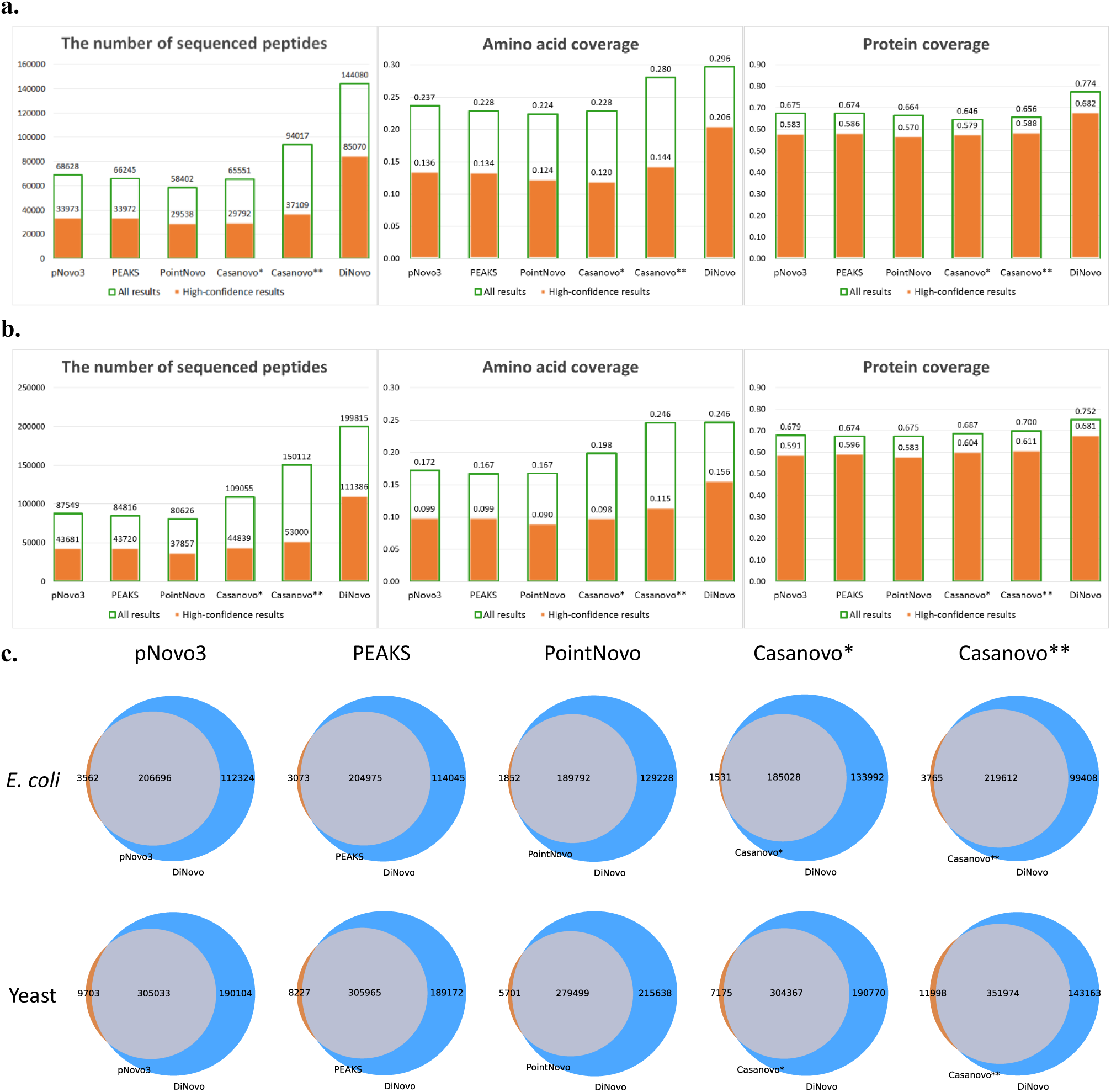
Comparison with different *de novo* sequencing algorithms. **(a)** *De novo* sequencing results on the *E. coli* datasets. **(b)** *De novo* sequencing results on the yeast datasets. Green hollow bars indicate all results, while orange solid bars indicate high-confidence (full fragment ion coverage) results. **(c)** Venn diagrams of high-confidence amino acids sequenced by DiNovo and other algorithms. Algorithms for comparison include pNovo3, PEAKS, PointNovo, Casanovo, and DiNovo. Casanovo* was trained on the 8-species dataset (excluding yeast), and Casanovo** was trained a large-scale dataset derived from the MassIVE Knowledge Base (MassIVE-KB). More detailed configurations of these algorithms are given in Methods.

The all results (green hollow bar) of DiNovo also surpassed those of other algorithms across the three levels for both *E. coli* and yeast datasets. More results are given in Supplementary Information (Figure S4-S6). All these results demonstrate that DiNovo is not only a high-coverage but also a high-confidence *de novo* peptide sequencing algorithm.

### Comparison with database search

As previously mentioned, the TD mapping method enables us to directly compare the results of *de novo* sequencing with those of database search at the same FDR level. To do this, we used the pFind search engine [27–29] to search the mass spectra against the protein sequence databases. The FDR of the database search results was controlled at 1%, and other search parameters are detailed in Methods. The results reported below were not filtered by high-confidence peptides.

As shown in Figure 6a and 6b, DiNovo sequenced much more peptides and identified much more spectra than pFind for all four proteases on the *E. coli* and yeast datasets. This advantage of DiNovo stems primarily from its use of complementarity of mirror spectra, which allows for the simultaneous accurate sequencing of mirror peptides that may be difficult to be sequenced alone. Of course, there is sequence redundancy in mirror peptides, and overall, DiNovo sequenced comparable numbers of amino acids and proteins to pFind. Importantly, there are significant overlaps between the results of DiNovo and pFind, demonstrating the reliability of both methods (Figure 6c, S7, S8). Specifically, DiNovo covered 85.8% (72.9%) of the amino acids and 92.4% (90.6%) of the proteins identified by pFind for the *E. coli* (yeast) datasets. Figure 6d further illustrates that the peptide sequences of spectra identified by both DiNovo and pFind are highly consistent. These results show that thanks to the mirror-protease strategy and the powerful DiNovo software, *de novo* sequencing is expected to become a practical and robust complement — or even an alternative — to traditional database search approach for peptide and protein identification. In this advancement, the TD mapping-based FDR control also plays a crucial role.

**Fig. 6.**
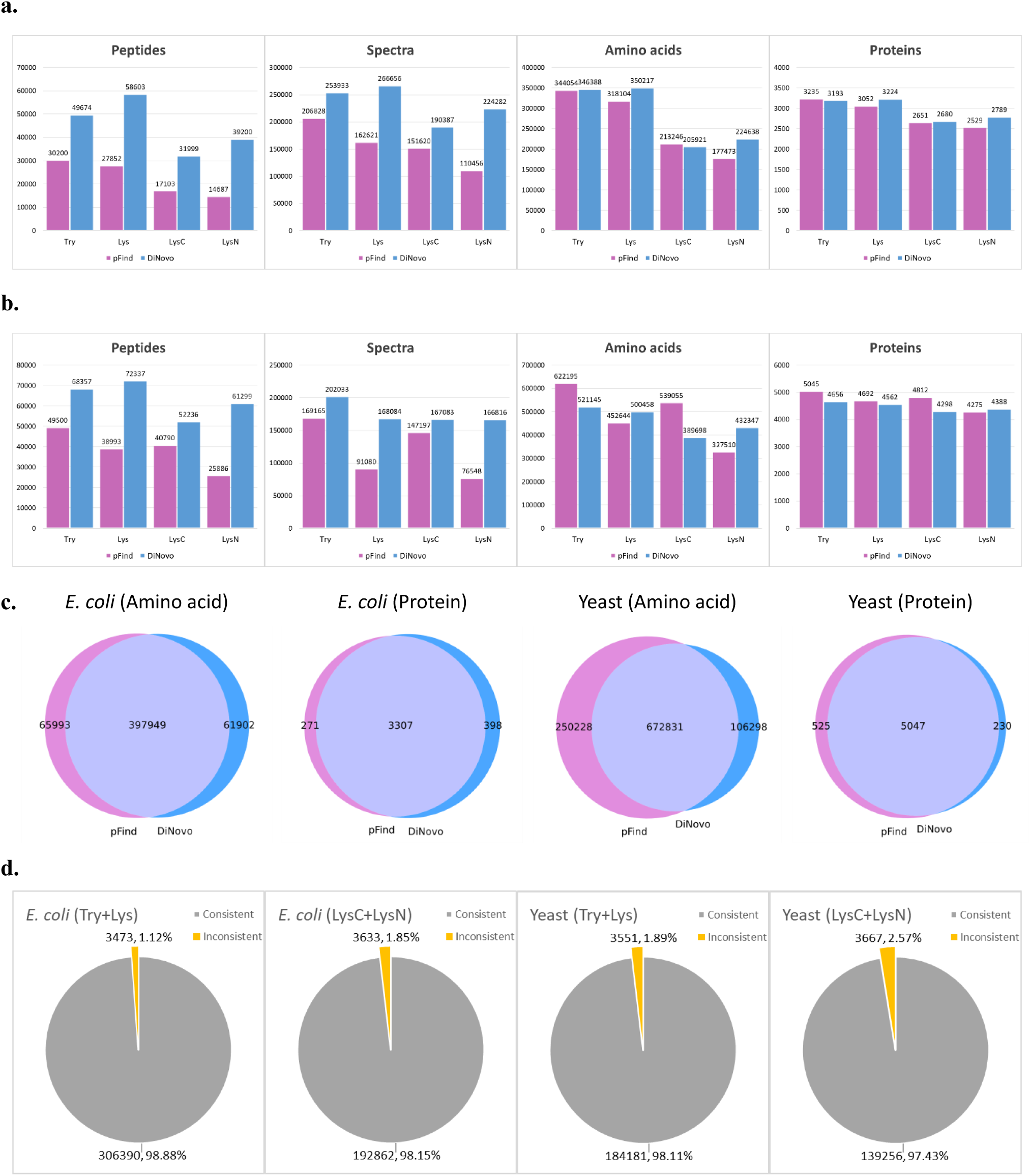
Comparison of *de novo* sequencing and database search results obtained by DiNovo and pFind. **(a)** *De novo* sequencing and database search results on the *E. coli* datasets. **(b)** *De novo* sequencing and database search results on the yeast datasets. **(c)** Venn diagrams of amino acids and proteins sequenced by DiNovo and pFind. **(d)** The consistency of peptide sequences of spectra identified by both DiNovo and pFind.

Interestingly, it can be observed that pFind showed proficiency in the identification of C-terminal digested peptides but was not as good as DiNovo at sequencing N-terminal digested peptides. This phenomenon suggests that current search engines, such as pFind, may have been optimized for C-terminal proteases, mainly trypsin, and can be improved for N-terminal proteases.

### Comparison of the two sequencing algorithms in DiNovo

There are two optional *de novo* peptide sequencing algorithms in DiNovo, i.e., MirrorNovo and pNovoM2. MirrorNovo is based on deep learning and runs on GPU, while pNovoM2 is based on graph theory and runs on CPU. We evaluated their sequencing accuracies and running speeds. Different mass thresholds were applied to control the FDR of each algorithm at 1%, as detailed in Table S2. As illustrated in Figure 7a and 7b, MirrorNovo sequenced more peptides than pNovoM2, but pNovoM2 had a much faster speed. Therefore, we recommend using MirrorNovo if one or more GPUs are available; otherwise, pNovoM2 is the better option for efficiency. Certainly, the two algorithms can be selected simultaneously, and their results can be combined by DiNovo to achieve optimal sequencing sensitivity.

**Fig. 7.**
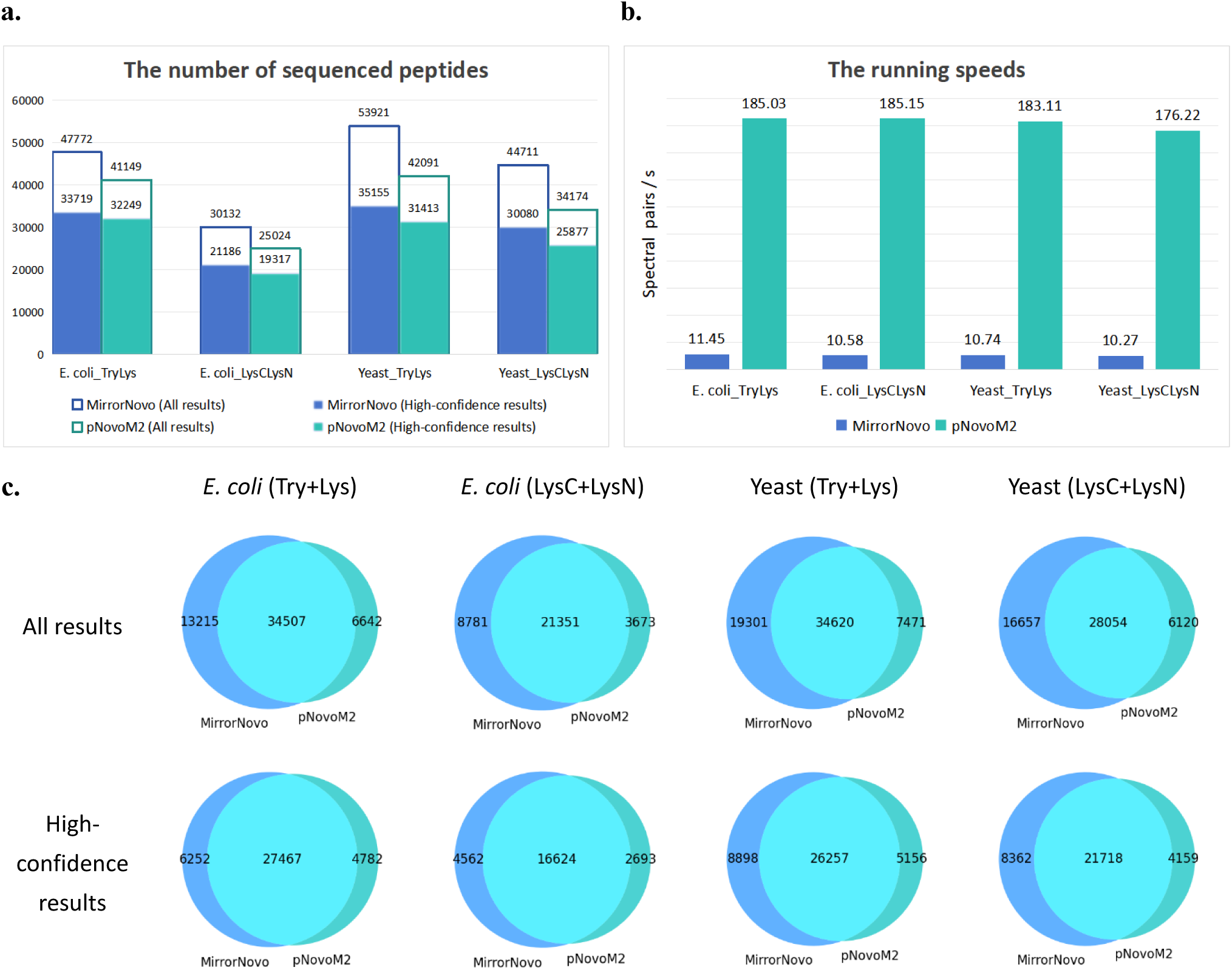
Comparison of MirrorNovo and pNovoM2. **(a)** The number of sequenced peptides. **(b)** The running speeds. **(c)** Venn diagrams of peptides sequenced by MirrorNovo and pNovoM2. MirrorNovo was run on GPU, while pNovoM2 was run on CPU. The *de novo* sequencing results are limited to mirror-protease sequencing due to the inability of pNovoM2 to perform single-protease sequencing.

Furthermore, we analyzed the intersection of the sequencing results of MirrorNovo and pNovoM2. As shown in Figure 7c, there was a significant overlap between the two algorithms, with each identifying a number of unique peptides. Finally, we analyzed the spectral pairs whose peptides were sequenced by both MirrorNovo and pNovoM2 (Figure S9). Under the 1% FDR, over 99% of these spectral pairs exhibited identical sequences, with a maximum of only 0.54% exhibiting differences, regardless of species and proteases. This outcome further demonstrated the reliability of the sequencing results and our quality control methodology.

## Discussion

In recent years, although deep learning has significantly advanced *de novo* peptide sequencing, the coverage and reliability of sequencing results remains unsatisfactory. This is largely due to the inherent limitations of current single protease-based LC-MS/MS experiments, i.e., incomplete fragment ions and undigested peptides. To address these challenges, we developed the DiNovo software that leverages multiple mirror proteases to enable higher-coverage and confidence *de novo* peptide sequencing. DiNovo is based on several advanced algorithms and, to our knowledge, is the first software suit supporting the full workflow of mirror-protease MS data analysis for *de novo* sequencing.

Experimental results demonstrate that the mirror-protease strategy employed in DiNovo can effectively improve fragment ion coverage of mass spectra. The average ion coverage of mirror spectra reached as high as 98%, much higher than separate spectra. Consequently, DiNovo achieved significantly higher sequencing accuracy than commonly used single-protease *de novo* sequencing algorithms such as PEAKS, pNovo3, PointNovo and Casanovo, which were applied to the separate spectra of four proteases. Specifically, DiNovo sequenced 110% to 194% more high-confidence peptides, which correspond to 36% to 74% higher amino acid coverage and 12% to 20% higher protein coverage. Remarkably, compared with the trypsin protease used alone, our using two pairs of mirror proteases led to 155% to 226% more high-confidence amino acids sequenced and 280% to 395% more high-confidence proteins identified.

Unlike current evaluation methods for *de novo* sequencing that rely on prior peptide identification by database search, DiNovo adopts TD database mapping to estimate the FDR of *de novo* peptides. TD mapping makes *de novo* sequencing a parallel method to database search for peptide identification with quality control. In our experiments, DiNovo identified comparable numbers of protein sequences to database search at the same FDR, and showed high degree of consistency. For the first time, we demonstrated the great potential of *de novo* sequencing as a practical alternative to database search. In this work, we ranked the *de novo* peptides by their precursor masses; however, an alternative and potentially better approach would be to rank them by a score, which could increase the number of peptides retained at the given FDR. Additionally, the pattern of mapping can be extended, e.g., from precise mapping to error-tolerant mapping to discover sequence variations or protein modifications. These are our ongoing work and will be reported elsewhere. Of course, general database-free quality control for *de novo* sequencing is still an open problem in the field with few attempts [30].

Recognition of mirror spectral pairs is a key step in mirror-protease *de novo* sequencing. In this paper we have proposed a new algorithm, named MirrorFinder, for this purpose. MirrorFinder directly compares two spectra and does not require pre-sequencing from separate spectra, and thus is less sensitive to spectral quality and is more efficient than pre-sequencing-based approach [23]. However, when the scale of spectra is very large, considering all spectral pairs can be a significant computational burden. Possible solutions to this problem are, for example, improving the precision of peptide precursor masses, using a tighter bound on retention time difference, or reducing data size by spectral clustering and combination.

MirrorNovo we developed is the first deep learning-based *de novo* sequencing algorithm for mirror spectra, and showed higher accuracy than traditional graph-based approach. As we know, the performance of DNNs greatly relies on the scale of training data, as illustrated by Casanovo [14,15] and most recent π-PrimeNovo [19], which were trained on ∼30 million spectra. Due to the limited availability of mirror spectra data, MirrorNovo has been trained on a relatively much smaller dataset in this paper. We can expect that as more training data becomes available in the future, there will be a lot of room for improvement in the performance of MirrorNovo.

Overall, it is reasonable to believe that the use of multiple mirror proteases can provide a promising solution for high-coverage and high-confidence *de novo* protein sequencing, and the DiNovo software we developed can serve as a powerful tool to facilitate mirror protease-based proteomics.

## Online Methods

### Protein sample preparation and LC-MS/MS analysis

Proteins of *E. coli* or yeast strains were extracted using a lysis buffer containing 8M urea, 150mM NaCl, 50 mM Tris-HCl (pH 7.5), 1mM phenylmethysulfonyl fluoride (PMSF), 1mM 2-chloroacetamide, and 1× cocktail (Roche, 11697498001). The proteins were reduced with 5mM DTT at 45℃ for 30min, and then fully alkylated with 15mM IAA at room temperature in the dark for 30min. Samples were processed by in-solution digestion or the FASP method as previously described, with slight modifications [31,32]. After reducing the urea concentration in the lysate to <1M by dilution or displacement, the proteins were aliquoted into four portions and digested with different proteases at an enzyme-to-protein ratio of 1:50. For trypsin/Lys-C or Lys-C alone digestion, the sample was solubilized in a buffer of 50mM ABC (pH 8.3), and trypsin and/or Lys-C were added for overnight digestion at 37℃. For LysargiNase/Lys-N digestion, the sample was solubilized in a buffer of 20mM HEPES, 10mM CaCl2, and 1mM Zn(Ac)2 (pH 8.3); after digestion with LysargiNase at 37℃ for 4h, Lys-N was added for overnight digestion. For Lys-N alone digestion, the sample was solubilized in a buffer of 50mM ABC and 1mM Zn(Ac)2 (pH 7.5-8.5), and Lys-N was added for overnight digestion at 37℃. All of trypsin, LysargiNase, Lys-C and Lys-N were provided by Enzyme & Spectrum (Beijing, China) [23, 33–35]. After digestion, 1% formic acid (FA) was added to stop the reaction. The digested peptides were desalted using an optimized StageTip [36].

LC-MS/MS analysis was conducted using an Orbitrap Q-Exactive HF mass spectrometer (Thermo Fisher Scientific, USA) coupled with an EASY-nLC 1000 system (Thermo Fisher Scientific, USA). The resulting peptides were resolved in buffer containing 1% acetonitrile and 1% FA, then loaded onto a 75μm I.D. ×20 cm LC column packed with 1.9 μm C_18_ reverse phase packing particles (Dr.Maisch GmbH, Germany). LC separation was performed with a 120min gradient from 6% to 45% buffer B (80% acetonitrile with 0.1% FA) at a flow rate of 300 nL/min. MS measurements were performed in data-dependent acquisition mode. For trypsin/Lys-C digested peptides, MS1 spectra were acquired with a survey scan (300-1400 m/z) at a resolution of 60,000 at 200 m/z, with an AGC target of 3e6 and a maximum injection time (MIT) of 30ms. Precursors with an intensity >3.3e4 were selected for MS2 analysis, collected at an MIT of 60ms or an AGC target of 5e4, then fragmented at a normalized collision energy (NCE) of 27. The resulting fragment ions were analyzed in the Orbitrap at a resolution of 15,000 at 400 m/z. For LysargiNase/Lys-N digested peptides, MS1 spectra were acquired with the same survey scan (300-1400 m/z) at a resolution of 60,000 at 200 m/z, an AGC target of 3e6 and an MIT of 50ms. Precursors with an intensity >1.3e4 were selected for MS2 analysis, collected with an MIT of 150ms or an AGC target of 1e5, then fragmented with an NCE of 27. The resulting fragment ions were analyzed in the Orbitrap at a resolution of 15,000 at 400 m/z.

### Datasets and preprocessing

The LC-MS/MS analysis of the *E. coli* and yeast proteins digested by four proteases resulted in eight datasets of MS2 spectra. The number of spectra in each dataset is shown in Table S3. In addition, we conducted database search using pFind v3.2.0 and calculated the pairing rate of the dataset at both spectrum and peptide levels (Table S3). For simplicity, we refer to trypsin/Lys-C mixed digestion as “trypsin” digestion, and LysargiNase/Lys-N mixed digestion as “LysargiNase” digestion.

Centroided spectra were extracted in Mascot generic format from RAW files using pParse [37]. In order to better recognize the mirror spectra, all mass spectra were preprocessed with the following steps. First, we performed two-step denoising, including global and local peak quantity control. For each spectrum, we retained the most abundant *N* peaks (*N* 200 by default), and then maintained the most abundant *K* peaks (*K* 4 by default) per mass bin (with a default bin width of 100Da). Additionally, we removed peaks corresponding to precursor ions and immonium ions. Second, we detected isotopic clusters and transformed each cluster into a singly charged monoisotopic peak with intensities combined. Third, to enhance the signals of backbone fragment ions, we accumulated the intensities from their isotopic and neutral-loss (-H2O and -NH3) peaks. It should be noted that these preprocessing steps were applied only to the mirror-spectra recognition process in DiNovo and not to the other algorithms in order to avoid any unexpected influence on their performance.

### Mirror-spectra recognition algorithm

In DiNovo, we developed a novel mirror-spectra recognition algorithm, named MirrorFinder, which is free of prior peptide sequencing from separate spectra as done in pNovoM [23]. The algorithm directly utilizes the information of precursor and fragment ions to recognize spectra generated from mirror peptides. To be specific, it first selects candidate spectral pairs whose precursor masses comply with the rule of mirror peptides (as outlined in Table S4) and whose retention time difference falls in a reasonable range. The rule states that each type of mirror peptides has a theoretical precursor mass difference and two theoretical fragment ion mass differences. Each candidate spectral pair is then assigned a matching score, reflecting the confidence level that they are indeed a pair of mirror spectra.

The matching score is calculated through the following steps (as illustrated in Figure 1b). First, a fixed mass range (-200 to +200 Da in this work) is divided into equal small intervals (e.g., 0.02 Da), with the total number of intervals denoted as *N*. Next, for the *i*-th interval, the number of fragment ion pairs whose mass differences fall in this interval is counted, and their intensities are summed, denoted as *count_i_* and *sum_inten_i_* (*i* 1, ⋯, *N*), respectively. Then, a statistic *s*_*i*_ is calculated for the *i*-th interval by multiplying the number of pairs by the summed intensities, which is

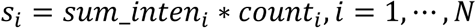

Finally, two *p*-values are computed for the intervals corresponding to two theoretical fragment ion mass differences (as shown in Table S4) from the distribution of the observed statistics. The smaller of the two *p*-values is denoted as *p*^∗^, and the matching score *S* is obtained by

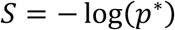

Obviously, a higher matching score indicates stronger evidence that the spectral pair is a pair of mirror spectra.

Subsequently, we use a target-decoy strategy [25,26] to filter spectral pairs according to their matching scores. In MirrorFinder, the matching score is closely related to the theoretical fragment ion mass differences, allowing us to directly use the score derived from a random mass difference as the decoy score to compete with the corresponding original (target) score. For each spectral pair, if the original score is higher than the decoy score, the spectral pair is labelled as a target spectral pair, and its final score is set as the original score. Otherwise, it is labelled as a decoy spectral pair, and its final score is set as the decoy score. Then the final scores are sorted in descending order, and the FDR of target spectral pairs with final scores above a given threshold *t* can be estimated as

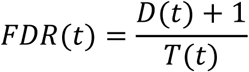

where *T*(*t*) and *D*(*t*) are the numbers of target and decoy spectral pairs, respectively. Through this strategy, we can filter target spectral pairs and control the FDR under a specified level, e.g., 2%. This approach eliminates the need to construct additional decoy spectra and offers a reasonable quality control standard for the recognized mirror spectra.

### DNN-based *de novo* sequencing algorithm for mirror spectra

We proposed a DNN-based *de novo* sequencing algorithm called MirrorNovo, which fully leverages complementary fragment ions in mirror spectra without the need to explicitly merge the spectra. In MirrorNovo, we preprocess each spectrum in a mirror spectral pair individually, padding the peak list with zeros if there are fewer than the top *N* peaks. To improve computational efficiency and reduce intensity variance, we convert the m/z values of all peaks to neutral masses and normalize their square-root-transformed intensities to the base peak.

As illustrated in Figure 1c, the input to the MirrorNovo model consists of two peak-ion matching matrices, each corresponding to one spectrum in a mirror spectral pair, along with the category of the spectral pair. We compute the two matrices by comparing the observed peak masses in each spectrum to 18 types of theoretical fragment ion masses (i.e., singly and doubly charged *b*, *y*, and *a* ions, their singly charged neutral-loss ions, and 6 types of internal ions), and then concatenate them along the peak dimension. Theoretical fragment ions are generated by enumerating all possible subsequent amino acids based on the already predicted prefix sequence. To ensure meaningful matrix concatenation, we align the two matrices according to the “homologous” fragmentation sites in the overlapping regions of the mirror peptides. For example, when iteratively predicting a trypsin-digested peptide of length *L* with either lysine or arginine (identified by MirrorFinder) at its C-terminal end, we align its *b_i_* (*y_L_*_-*i*_) fragments with the *b_i_*_+1_ (*y_L_*_-*i*-1_) fragments derived from the LysargiNase-digested peptide, which has either lysine or arginine at its N-terminal end.

MirrorNovo employs a T-Net architecture [13] with three one-dimensional convolutional layers to extract peak features from the concatenated matching matrices. A subsequent GRU layer captures both short- and long-term dependencies across peaks while maintaining the dimensionality of the input and output matrices. Next, three fully connected (FC) layers are utilized to learn higher-level feature representations, with batch normalization applied between them to enhance training stability and accelerate convergence. The Rectified Linear Unit (ReLU) activation function introduces non-linearity to prevent the collapse of adjacent linear layers. Finally, a softmax layer outputs probabilities for 20 amino acids, where isoleucine and leucine are not distinguished.

MirrorNovo predicts complete peptide sequences from mirror spectral pairs through iterative amino acid inference. At each iteration, it employs a beam search strategy to avoid local optima and utilizes a knapsack algorithm to reduce the number of candidate amino acids. A bidirectional prediction strategy is integrated to mitigate the loss of correct sequences. The iteration terminates when the difference between the prefix mass and precursor mass falls within 20 ppm. The top *M* (*M* 10 by default) candidate peptides are retained based on peptide probability scores, calculated as the average of individual amino acid probabilities. To enhance data utilization, DiNovo also includes a single-protease *de novo* sequencing algorithm, Denovo-GCN [18], which allows peptide sequencing from spectra that cannot form mirror spectral pairs.

In this study, MirrorNovo was trained on three high-resolution tandem mass spectrometry datasets from protein samples of Vero, MC2155, and human testis. Annotated mirror spectral pairs were constructed based on pFind (v3.2.0) search results at a 1% FDR. Since a pair of mirror peptides may correspond to multiple mirror spectral pairs, and to ensure high-quality PSMs and sequence diversity, we ranked the mirror spectral pairs by pFind’s final scores and selected up to the top *K* highest-scoring pairs. Only spectral pairs of types A, B, and C with charge states of 2+ or 3+ were retained for the final annotated dataset. The dataset was split into training and validation sets at a 9:1 ratio based on peptide sequences (details in Table S5), ensuring no sequence overlap between the sets. The single-protease sequencing model, Denovo-GCN, was trained on all trypsin- and LysargiNase-digested PSMs from the same dataset.

### Graph-based *de novo* sequencing algorithm for mirror spectra

In addition to the DNN-based sequencing algorithm MirrorNovo, we also developed a graph-based sequencing algorithm, pNovoM2, which generates candidate peptides through spectrum graphs. Building on pNovoM [23], we have enhanced the performance of peptide sequencing and expanded the types of mirror peptides that can be sequenced, enabling pNovoM2 to deliver more results than pNovoM. Furthermore, we have optimized the speed and memory management of pNovoM2, facilitating rapid *de novo* peptide sequencing on large-scale datasets without the need for GPU support.

Figure S10 illustrates the schematic diagram of pNovoM2. Unlike pNovoM, pNovoM2 interprets not only the merged mirror spectra but also the separate spectra. This strategy allows pNovoM2 to sequence peptides even when MirrorFinder fails to find the mirror spectral pairs. In the workflow of pNovoM2, all spectra are first preprocessed, with each generating a spectrum graph similar to pNovo [8]. If MirrorFinder recognizes the corresponding mirror spectrum for the input spectrum, the spectrum graphs of the two spectra will be merged and used for subsequent analysis. Otherwise, the spectrum graph of the input spectrum will be used directly. For the merged spectrum graph, an edge is created between two nodes if their mass difference matches the total mass of any two or three amino acids. However, for the spectrum graph from a separate spectrum, the maximum number of amino acids is increased to five, due to the relatively lower abundance of signal peaks. Our scoring function aligns with that of pNovo, but we’ve made slight adjustments to better accommodate the merged spectrum graph. If the mass of a node in one spectrum graph differs from the mass of a node in another spectrum graph by a specific value, such as the mass of the amino acid K or R in the case of trypsin and LysargiNase, the peaks corresponding to these nodes are likely signal peaks of fragment ions. These nodes are then assigned a higher score. The pDag [38] algorithm is subsequently used to identify the top *N* highest-scoring paths from the source node to the target node. For the merged spectrum graph, the value of *N* is set to 20; for the spectrum graph from a separate spectrum, it is set to 40. All peptides corresponding to these paths from the spectrum graphs are rescored together, and a final list of candidate peptides is generated.

### *De novo* sequencing results evaluation

In this paper, we invented a simple yet effective method for evaluating *de novo* sequencing results, called TD mapping (as illustrated in Figure 2a). This method requires the existence of sequence database of proteins to be sequenced, but eliminates the need for peptide identification by traditional database search. Through the introduction of decoy database, it addresses the lack of quality control in existing mapping-based methods [16].

Building on the TD method [25,26], we first generate a decoy protein database by simply reversing or shuffling the sequences in the original database. Then, we map all the *de novo* sequences separately to the protein sequences in the target and decoy databases. Those successfully mapped peptides are retained and subjected to subsequent analysis. Unmatched peptides are excluded, as well as those mapped to both target and decoy sequences. Next, mapped peptides are ordered according to their masses, and the FDR of target peptides with masses above a threshold *m* is estimated as

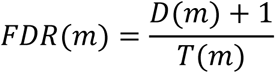

where *T*(*m*) and *D*(*m*) are the numbers of target and decoy peptides, respectively. Finally, an appropriate mass threshold is selected so that the FDR is controlled under the desired level.

### Configuration of *de novo* sequencing and database search

To fairly compare the performance of *de novo* sequencing software tools, we configured different tools with the same parameters. Both precursor and fragment tolerances were set to 20 ppm. Carbamidomethyl of cysteine (C) was specified as a fixed modification and oxidation of methionine (M) was considered as a variable modification. In addition, to perform a horizontal comparison between *de novo* sequencing and database search, all raw files were analyzed using pFind v3.2.0 for peptide identification. The detailed configurations for each software tool are as follows:

- λ **DiNovo**: We offer two optional *de novo* sequencing algorithms in DiNovo, i.e., MirrorNovo and pNovoM2. By default, MirrorNovo is used for optimal performance, unless otherwise specified. The MirrorNovo model is trained with the following parameters: optimizer = Adam, loss function = Focal, maximum epochs = 5, batch size = 20, learning rate = 1×10^-3^, decay factor = 0.5. When comparing these two algorithms (as discussed in Section 2.7), we set the number of processes of pNovoM2 as the batch size (9) of MirrorNovo to ensure consistency in the number of parallel operations. Additionally, MirrorNovo was trained on an NVIDIA GeForce RTX 3090 GPU, while pNovoM2 was executed on an Intel(R) Xeon(R) Silver 4214R CPU.
- λ **PEAKS**: We acquired the commercial software Peaks-Online 11, and used its built-in *de novo* peptide sequencing algorithm to analyze *E. coli* and yeast datasets in Sections 2.4 and 2.5. Peaks-Online 11 utilizes an integrated deep learning-based *de novo* peptide sequencing algorithm (PointNovo [13]). However, the specifics of the training set used to train the model remain undisclosed to us.
- λ **pNovo3**: pNovo3 [12] uses the pDag [38] algorithm to find the maximum scoring path within the spectrum graph, after which it rescores candidate peptides using an SVM model. The peptide with the highest score is then reported as the output. For our analysis in Section 2.5, we used the latest version of pNovo3, pNovo v3.1.5, which was released on December 6, 2023.
- λ **PointNovo**: PointNovo [13] is a neural network model that employs a T-Net structure, specifically designed for order-invariant data. This design allows it to be applicable to mass spectra of varying resolutions without increasing computational complexity. Since the authors of PointNovo did not provide a trained model file, and the original 9-species training dataset used in [13] included the yeast species, we excluded the yeast species and trained a PointNovo model from scratch on the remaining 8-species dataset for our analysis.
- λ **Casanovo* and Casanovo**:** Casanovo [14,15] is a deep learning algorithm that employs a transformer neural network architecture to translate the sequence of peaks in a tandem mass spectrum into the corresponding amino acid sequence of the generating peptide. Notably, Casanovo does not constrain the peptides it generates by precursor mass, so the precursor masses of its generated peptides may not be equal to those provided by the spectra. In contrast, all the other algorithms mentioned above use precursor mass as a constraint for *de novo* peptides. Therefore, it would be unfair to Casanovo to directly use the highest-scoring results without considering the precursor mass error. However, if we filter out peptides generated by Casanovo with precursor mass errors exceeding 20 ppm, some low-quality spectra will not yield any sequencing results. As a result, these spectra will be discarded in the results of Casanovo, and the TD mapping method will be applied only to the remaining spectra with higher quality. This could unfairly underestimate the FDR of Casanovo compared to other software tools. For a fairer comparison, we adopted a compromise strategy: we filtered out peptides generated by Casanovo with precursor mass errors exceeding 20 ppm and applied the same peptide mass thresholds to the Casanovo results as we did to DiNovo. For our analysis in Section 2.5, we used two models of Casanovo 4.2.0, which was released on May 14, 2024. The first model, referred to as Casanovo*, was trained by us from scratch on the 8-species dataset, similar to the approach used with PointNovo. The second model, referred to as Casanovo**, is the latest version provided by its authors, trained on a large-scale dataset derived from the MassIVE Knowledge Base (MassIVE-KB), which consists of 30 million PSMs. It is important to note that Casanovo** is a trained model that does not allow for the specification of modifications. For a fair comparison, we selected the highest-ranking peptide without unexpected modifications as the result for each spectrum.
- λ **pFind:** The *E. coli* dataset, released in August 2015 on UniProt, includes 4,319 protein entries along with contaminant protein sequences. The yeast dataset, released in January 2015 on UniProt, contains 6,726 protein entries, also with contaminant protein sequences. For the close search parameters, both precursor and fragment mass tolerances were set to 20 ppm. Carbamidomethyl of cysteine (C) was specified as a fixed modification, whereas oxidation of methionine (M) was set as a variable modification. The maximum number of modification sites per peptide sequence was limited to 3. Upon completing the database search, peptides and proteins were filtered using a 1% FDR at both the spectrum and protein levels.

### Design of DiNovo software

Previously, the only software available for mirror-protease *de novo* sequencing is pNovoM, which requires users to manually carry out step-by-step operations for pre-sequencing, mirror-spectra recognition, and *de novo* sequencing (note that the programs for the first two steps were not publicly released). This makes the use of it cumbersome and not user-friendly. As a result, there remains a need for software that can recognize and sequence mirror spectra with a simple, one-click operation.

To meet this need, we integrate the various algorithms developed in this paper into a complete software solution named DiNovo, making it easily accessible to participants in the field. The DiNovo software includes the following modules: configuration settings, MS data I/O, spectra preprocessing, mirror-spectra recognition, *de novo* sequencing, and real-time log output during the process. As illustrated in Figure S11, parallelization is fully supported in each analytical component of DiNovo, significantly improving computational efficiency and reducing overall processing time.

In DiNovo, error-tolerant precursor mass indexes are built for all spectra to enable high-speed mirror-spectra recognition. As a result, the time complexity of DiNovo for selecting candidate spectral pairs involving *n* trypsin-digested spectra and *m* LysargiNase-digested spectra is *O*(*m + n*), in contrast to *O*(*mn*) by brute force. For example, DiNovo takes approximately 20 seconds to select candidate spectral pairs from a pool of 10 million spectral pairs, 1,000 times faster than brute force. On the 16 million spectral pairs of *E. coli* dataset, DiNovo performed mirror-spectra recognition 2.1 and 46.5 times faster than pMerge (the mirror-spectra recognition module of pNovoM, unreleased) with 1 and 30 processes on a workstation, respectively. In combination of mirror-spectra recognition and sequencing, DiNovo completed the whole task four to five times faster than pMerge and pNovoM.

## Acknowledgements

This work is supported by the National Key R&D Program of China (Grants 2022YFA1304600 and 2022YFA1004801), the National Natural Science Foundation of China (Grant 32070668), and the CAMS Innovation Fund for Medical Sciences (2019-I2M-5-017 and 2022-I2M-CandT-B-082).

## Supplementary Information

### List of Supplementary Figures

**Supplementary Figure 1.**
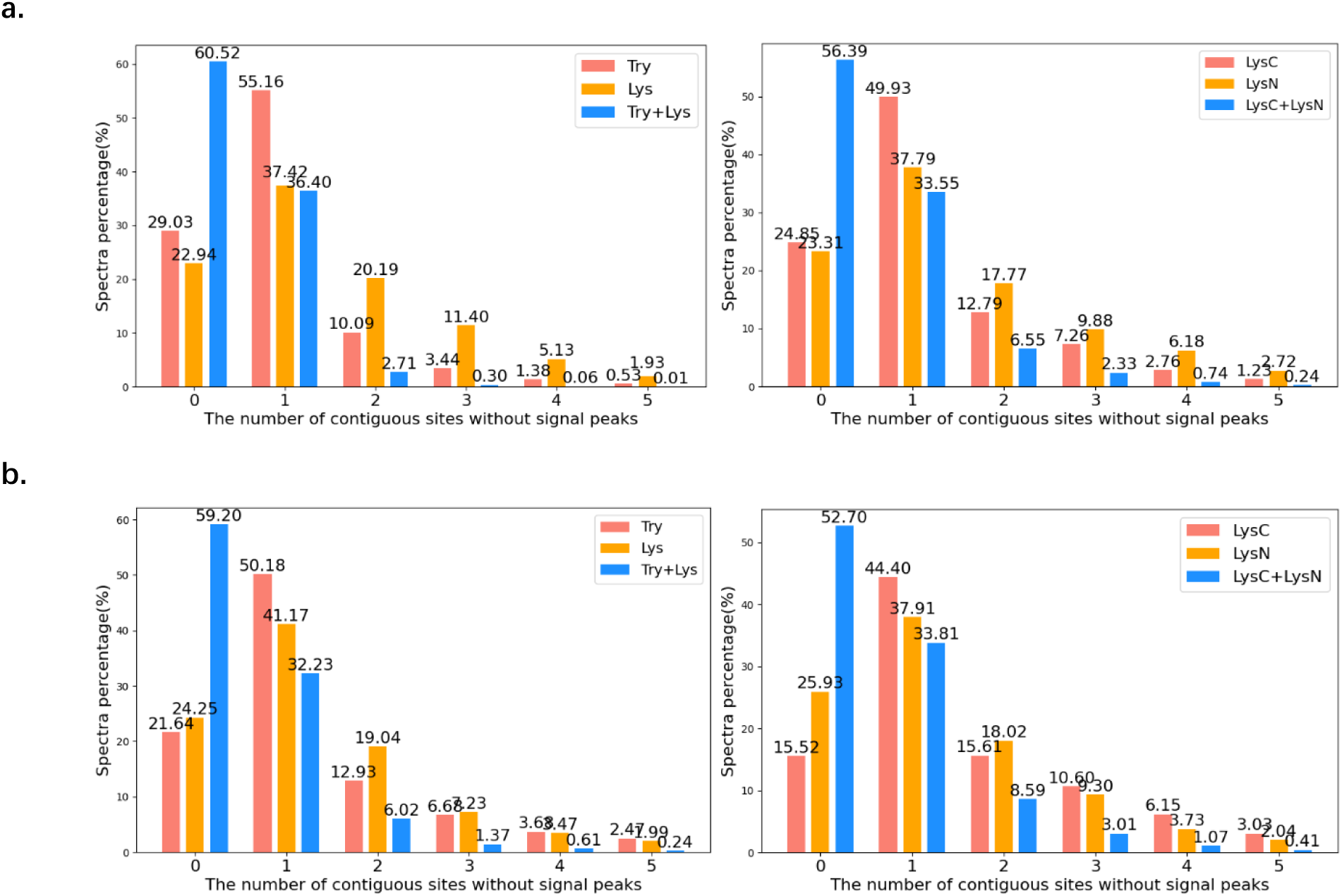
The distribution of the number of contiguous sites without signal peaks. (a) *E. coli* datasets. (b) Yeast datasets. Try and Lys are abbreviations of trypsin and LysargiNase, respectively.

**Supplementary Figure 2.**
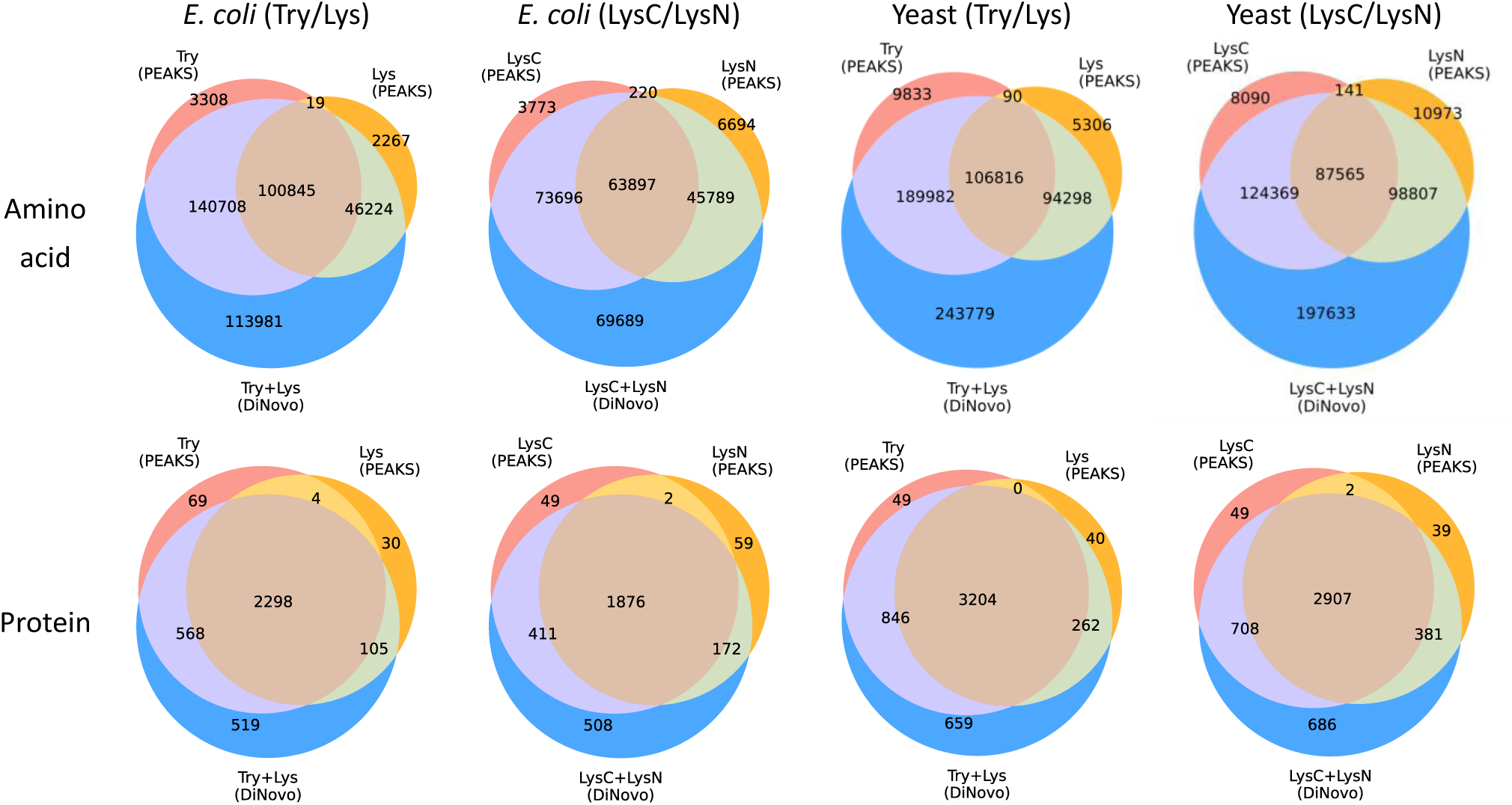
Venn diagrams of amino acids and proteins sequenced by DiNovo and PEAKS. These results are not filtered by high-confidence (full fragment ion coverage) peptides.

**Supplementary Figure 3.**
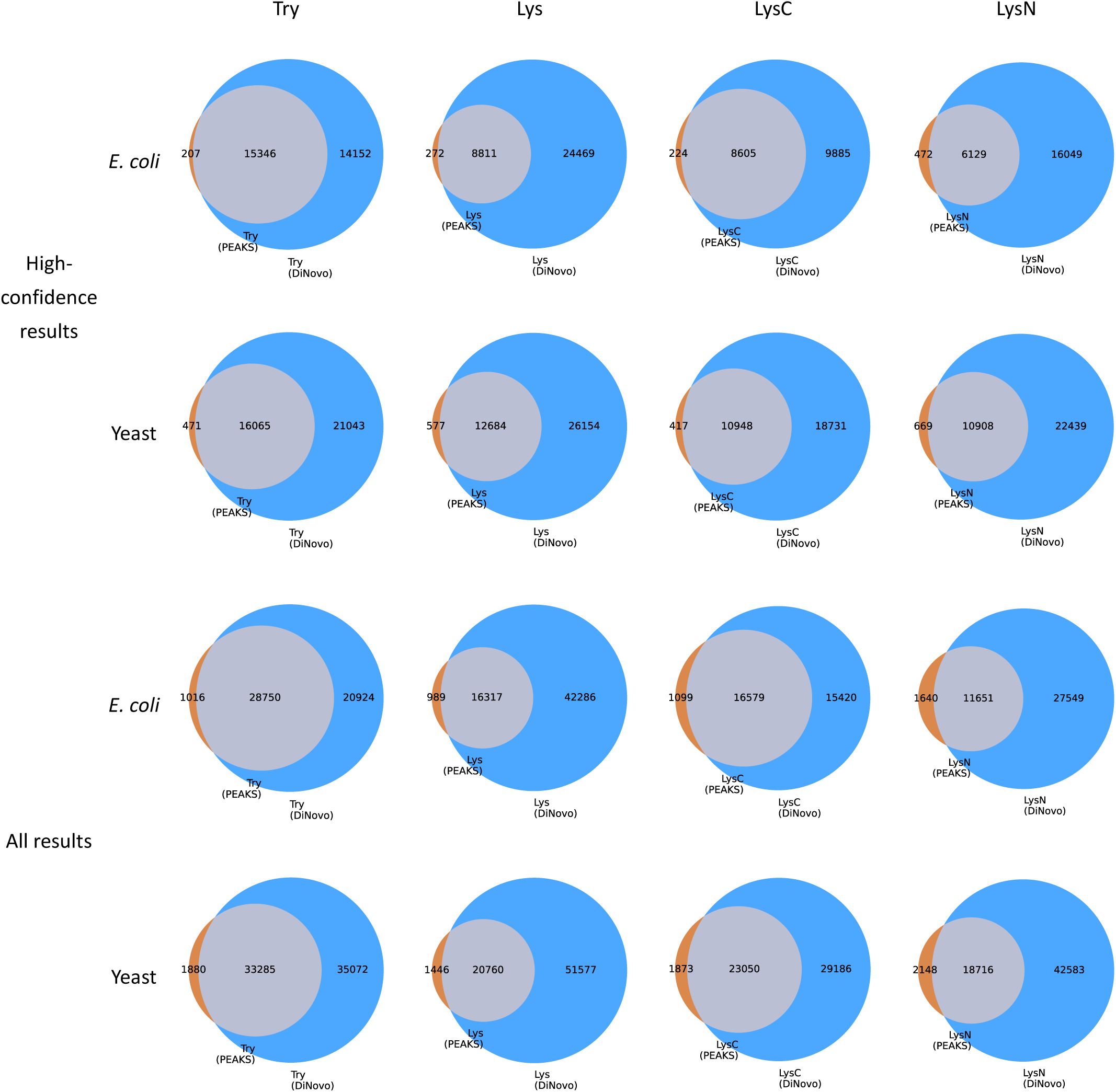
Venn diagrams of peptides sequenced by DiNovo and PEAKS.

**Supplementary Figure 4.**
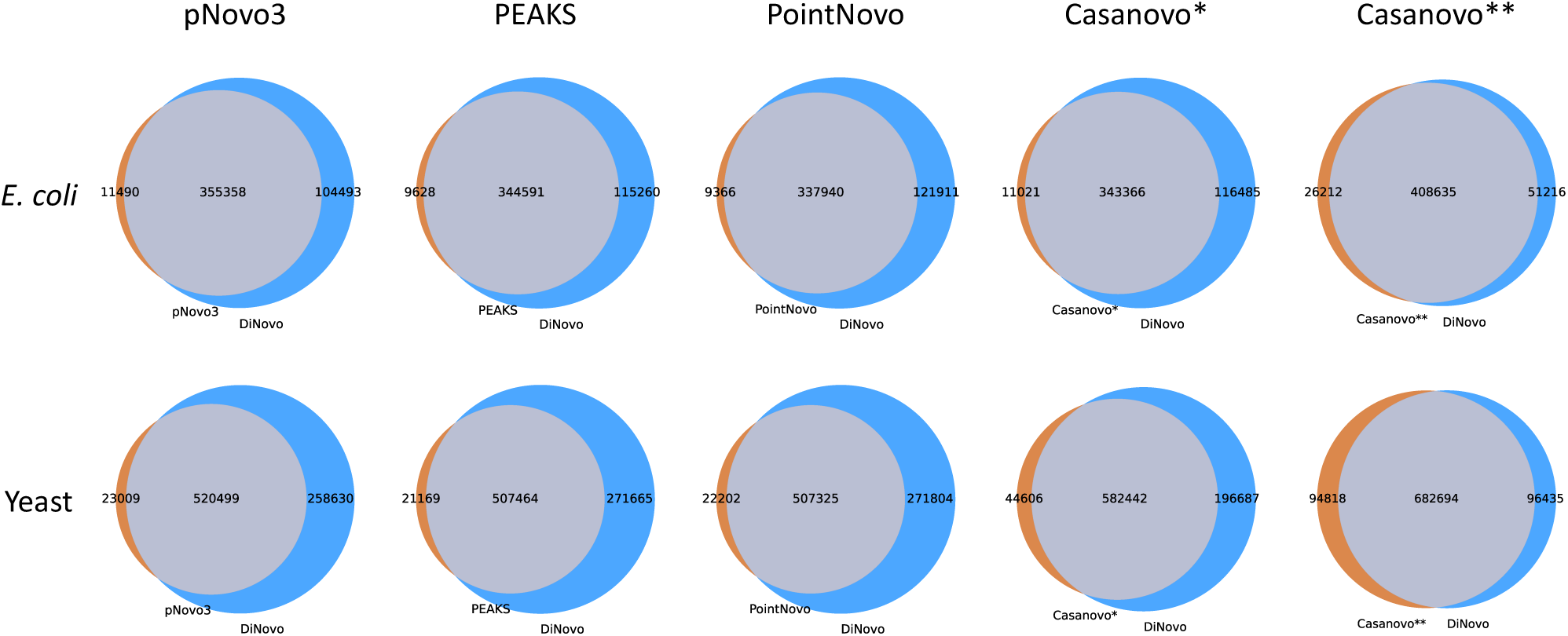
Venn diagrams of amino acids sequenced by DiNovo and other algorithms. These results are not filtered by high-confidence (full fragment ion coverage) peptides. Algorithms for comparison include pNovo3, PEAKS, PointNovo, Casanovo, and DiNovo. Casanovo* was trained on the 8-species dataset (excluding yeast), and Casanovo** was trained a large-scale dataset derived from the MassIVE Knowledge Base (MassIVE-KB).

**Supplementary Figure 5.**
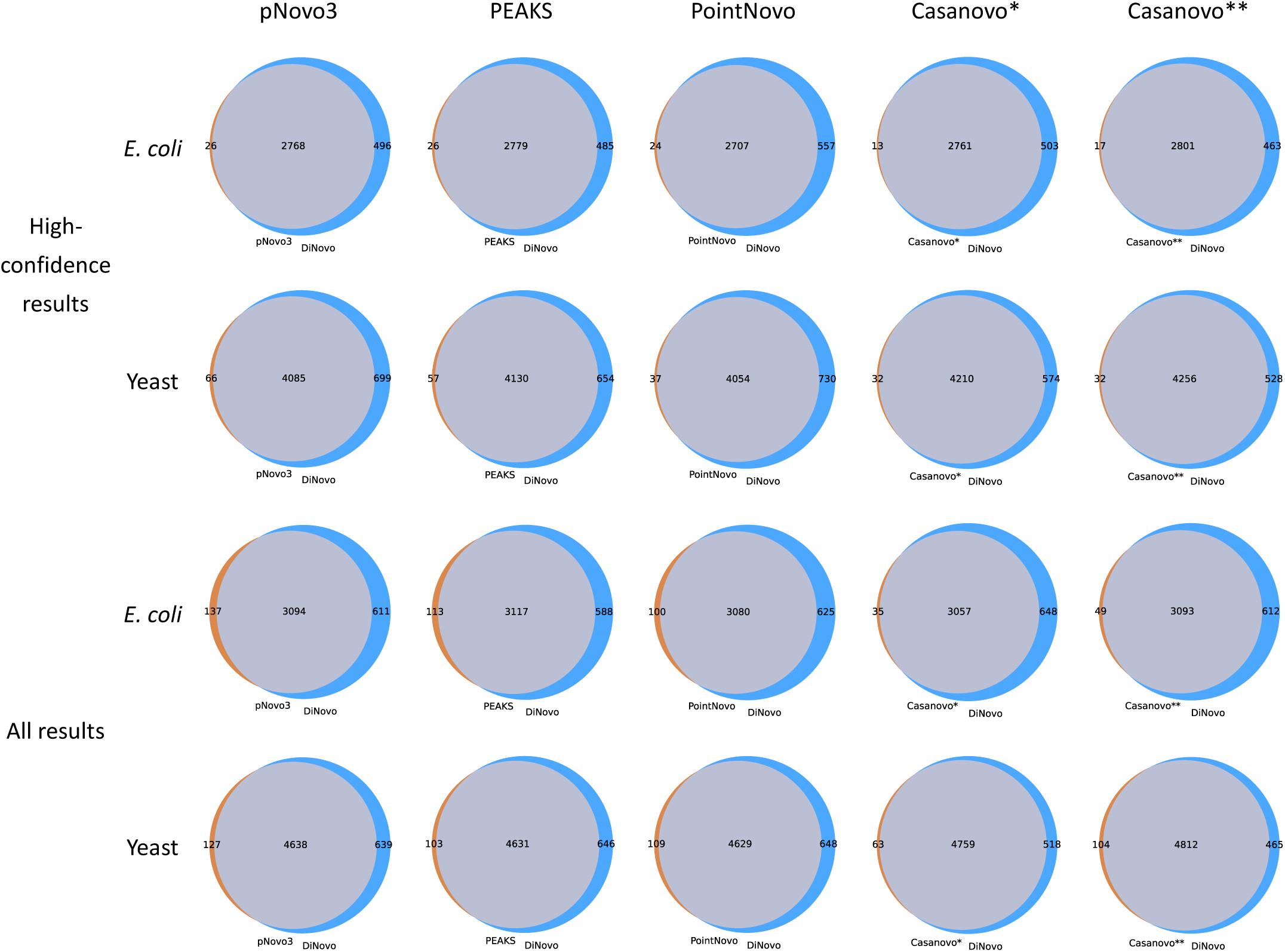
Venn diagrams of proteins sequenced by DiNovo and other algorithms.

**Supplementary Figure 6.**
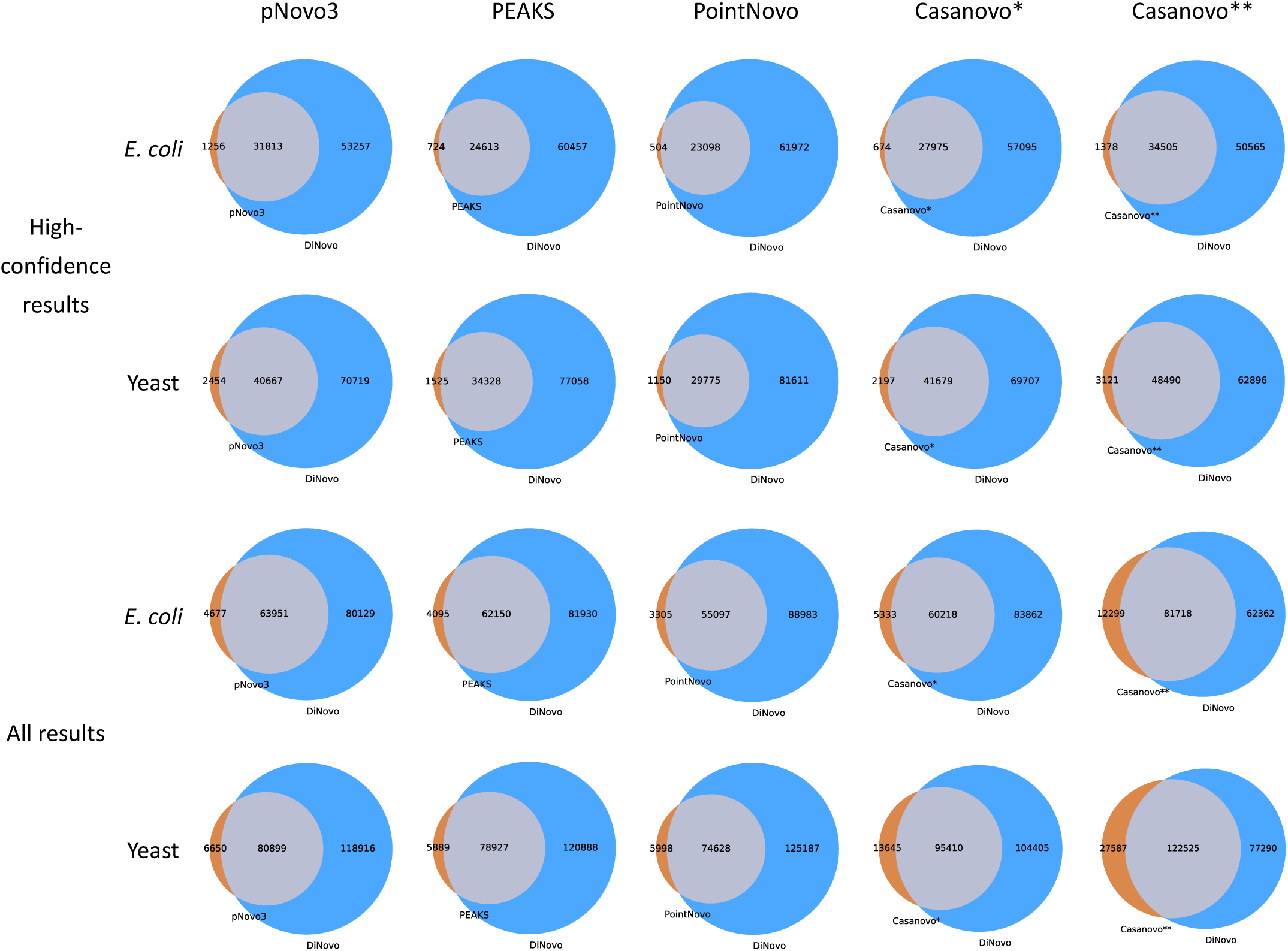
Venn diagrams of peptides sequenced by DiNovo and other algorithms.

**Supplementary Figure 7.**
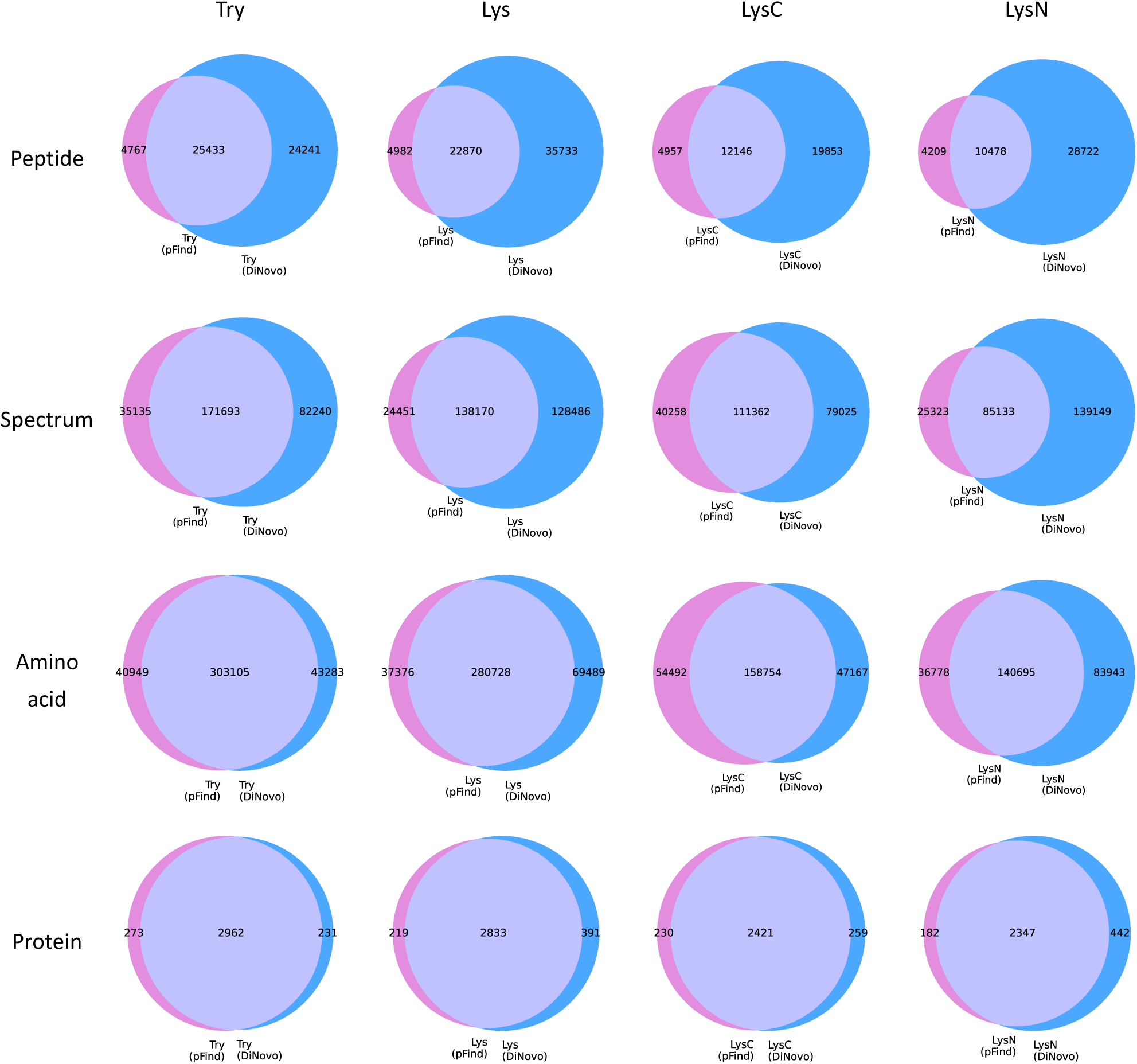
Venn diagrams of peptides, spectra, amino acids and proteins sequenced by DiNovo and pFind on the *E. coli* datasets.

**Supplementary Figure 8.**
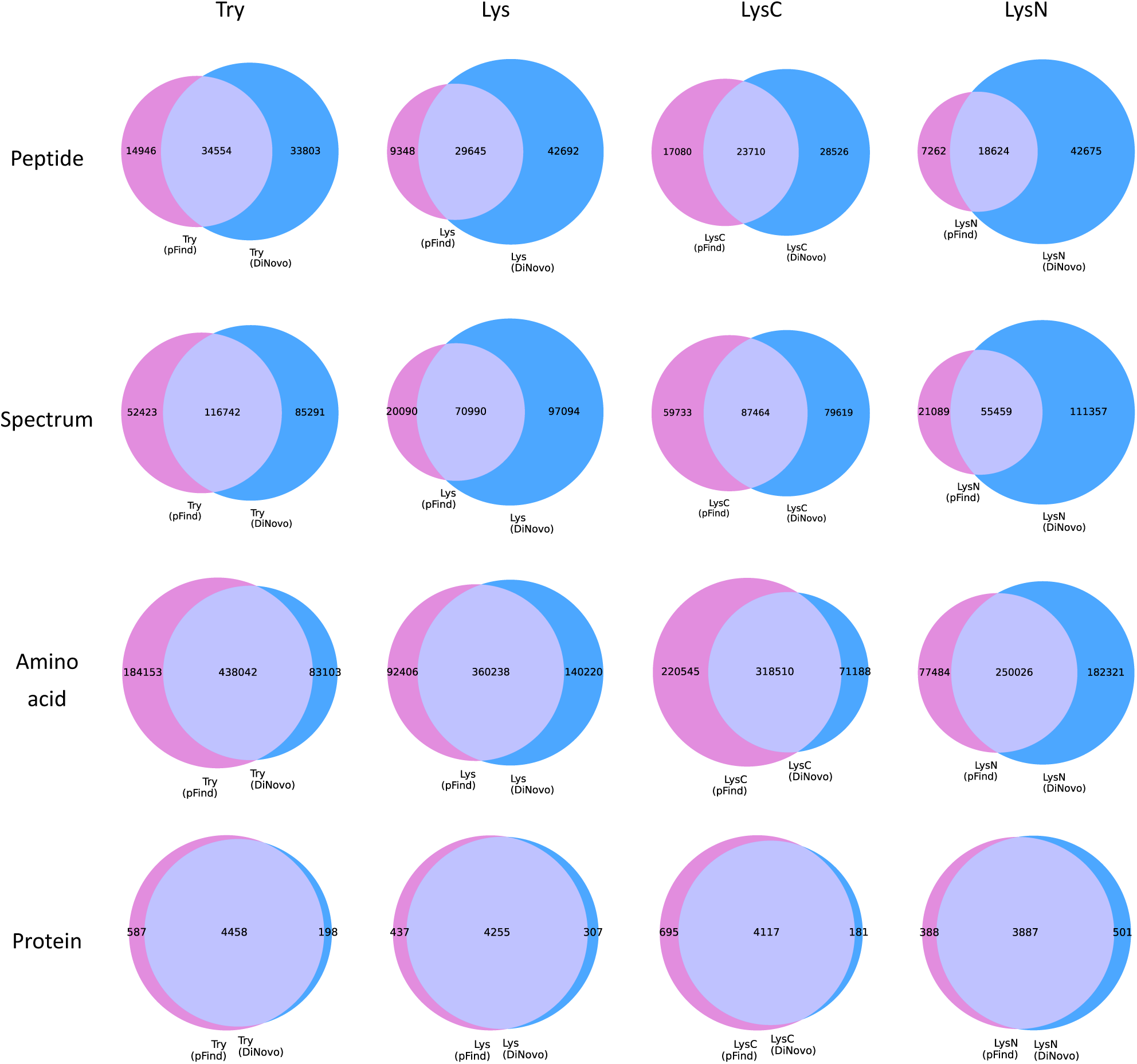
Venn diagrams of peptides, spectra, amino acids and proteins sequenced by DiNovo and pFind on the yeast datasets.

**Supplementary Figure 9.**
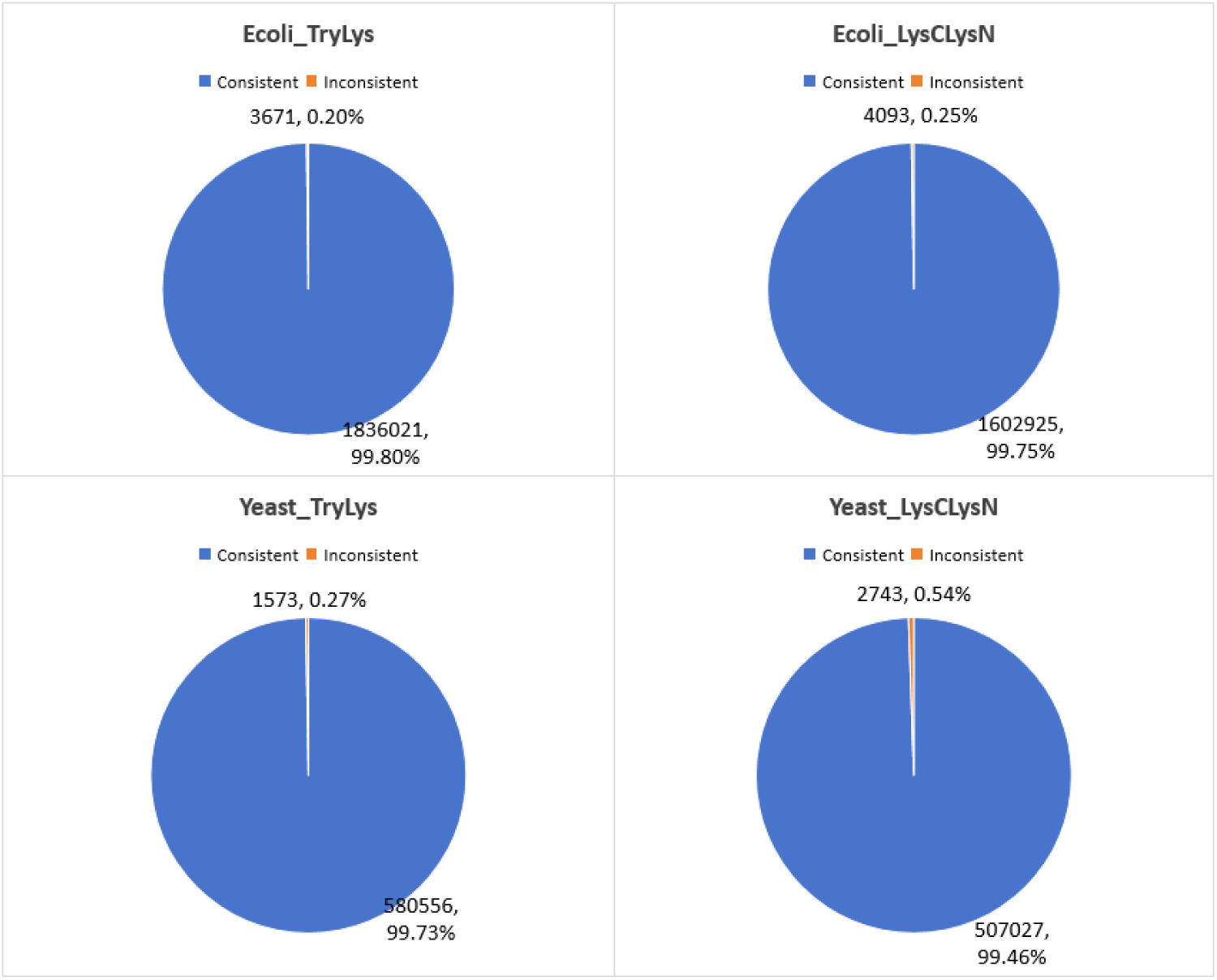
Spectral pairs whose peptides sequenced by both MirrorNovo and pNovoM2. The blue color represents that the MirrorNovo sequence is consistent with the pNovoM2 sequence, otherwise it is represented by orange.

**Supplementary Figure 10.**
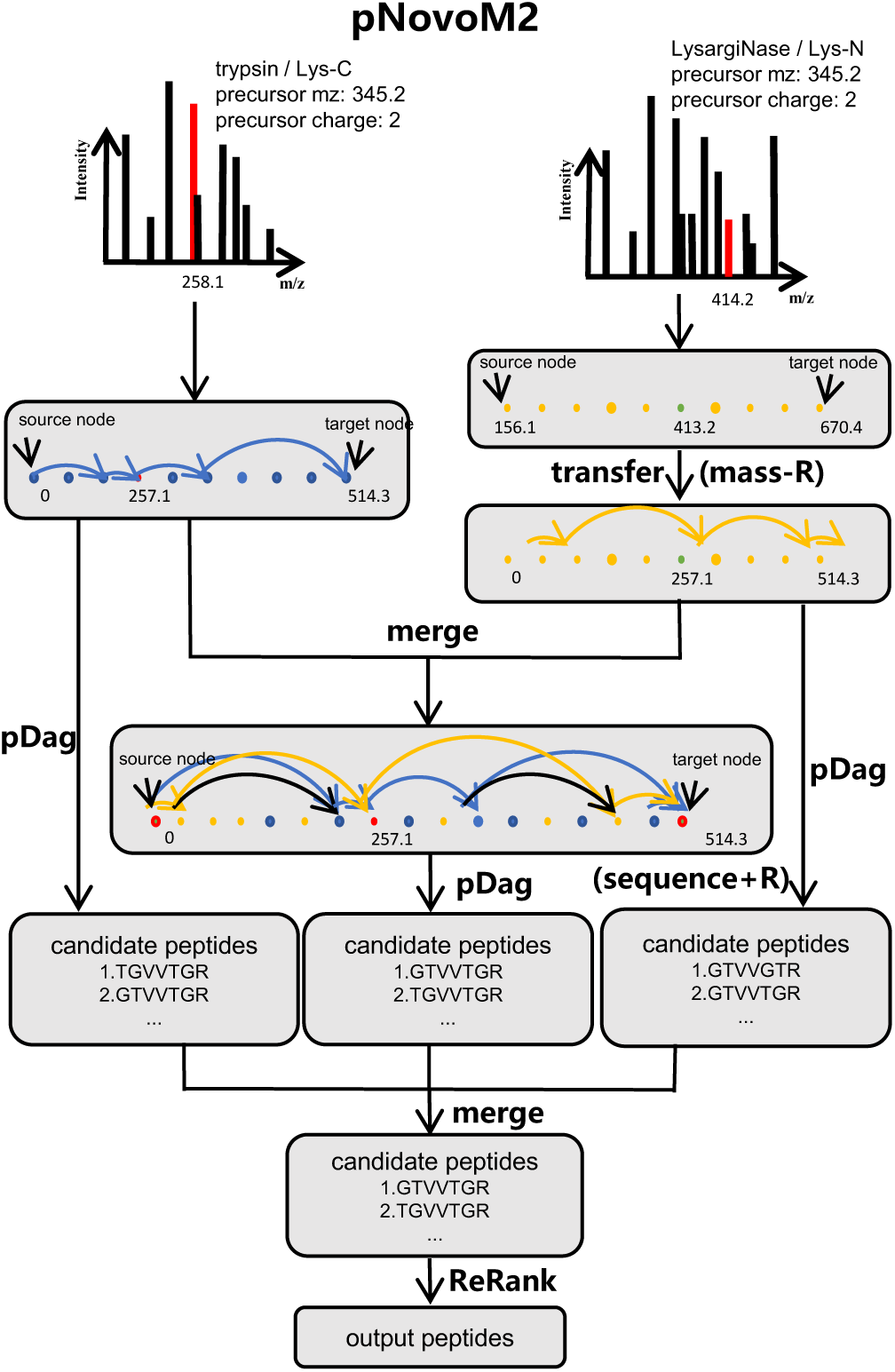
The schematic diagram of pNovoM2.

**Supplementary Figure 11.**
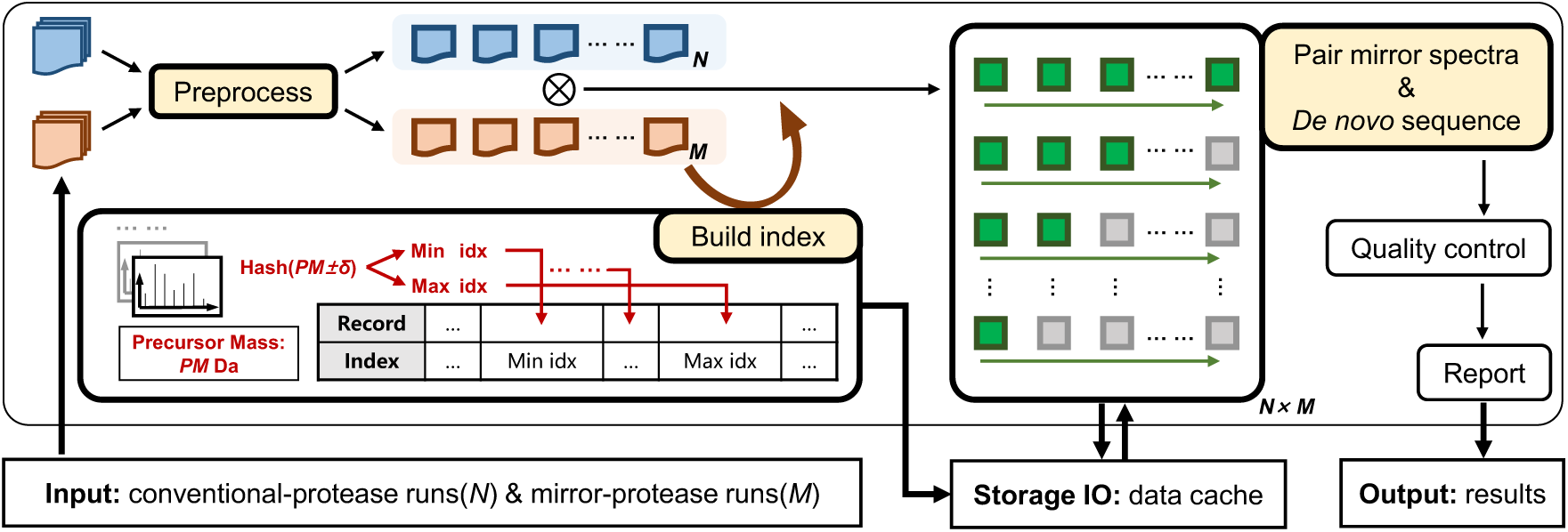
Design of DiNovo software.

### List of Supplementary Tables

**Supplementary Table 1.** Mass threshold for quality control used by different software tools (Dalton)

|  | pNovo3 | PEAKS | PointNovo | Casanovo | DiNovo |
| --- | --- | --- | --- | --- | --- |
| *E. coli*: Try/Lys | 720/770 | 750/790 | 760/830 | 740/740 | 740 |
| *E. coli*: LysC/LysN | 730/760 | 760/800 | 780/850 | 760/760 | 760 |
| Yeast: Try/Lys | 730/770 | 790/820 | 790/850 | 770/770 | 770 |
| Yeast: LysC/LysN | 750/780 | 790/820 | 810/860 | 770/770 | 770 |

**Supplementary Table 2.** Mass threshold for quality control used by MirrorNovo and pNovoM2 (Dalton)

|  | MirrorNovo | pNovoM2 |
| --- | --- | --- |
| *E. coli*: Try+Lys | 660 | 710 |
| *E. coli*: LysC+LysN | 670 | 710 |
| Yeast: Try+Lys | 760 | 780 |
| Yeast: LysC+LysN | 740 | 750 |

**Supplementary Table 3.** The size and pairing rate of the datasets.

|  |  | Number of spectra | Spectrum pairing rate | Peptide pairing rate |
| --- | --- | --- | --- | --- |
| *E. coli* | Trypsin | 1,378,808 | 84.32% | 67.74% |
|  | LysargiNase | 1,364,202 | 87.73% | 75.09% |
|  | Lys-C | 1,829,193 | 82.41% | 61.68% |
|  | Lys-N | 1,630,987 | 90.59% | 78.19% |
| Yeast | Trypsin | 1,111,519 | 68.06% | 48.52% |
|  | LysargiNase | 725,829 | 76.96% | 64.08% |
|  | Lys-C | 1,163,099 | 64.90% | 42.79% |
|  | Lys-N | 873,543 | 82.47% | 70.52% |

**Supplementary Table 4.** The rule of mirror peptides.

| Category | Mirror peptides |  | Theoretical precursor mass difference | Theoretical fragment ion mass difference |
| --- | --- | --- | --- | --- |
| A | PEPTIDER | RPEPTIDE | 0 Da | -156 Da, 156 Da |
|  | PEPTIDEK | KPEPTIDE | 0 Da | -128 Da, 128 Da |
| B | PEPTIDER | KPEPTIDE | 28 Da | -128 Da, 156 Da |
| C | PEPTIDEK | RPEPTIDE | -28 Da | -156 Da, 128 Da |
| D | PEPTIDEK | PEPTIDE | 128 Da | 0 Da, 128 Da |
| E | PEPTIDER | PEPTIDE | 156 Da | 0 Da, 156 Da |
| F | PEPTIDE | KPEPTIDE | -128 Da | -128 Da, 0 Da |
| G | PEPTIDE | RPEPTIDE | -156 Da | -156 Da, 0 Da |

**Supplementary Table 5.** The training and valid datasets of MirrorNovo and Denovo-GCN.

| Species | MirrorNovo |  | Denovo-GCN |  |
| --- | --- | --- | --- | --- |
|  | Training Dataset<br>(PSM pairs) | Valid Dataset<br>(PSM pairs) | Training Dataset<br>(PSMs) | Valid Dataset<br>(PSMs) |
| Testis | 279241 | 31027 | 347233 | 40302 |
| Vero | 74318 | 8258 | 80922 | 9720 |
| MC2155 | 120889 | 13433 | 122481 | 15335 |

